# Timed STING Inhibition Mitigates Gastrointestinal GvHD While Preserving Graft-versus-Leukemia Activity After Allo-HSCT

**DOI:** 10.64898/2026.07.17.738478

**Authors:** Chiara Suriano, Omer Khalid, Paul Heinrich, Kaiji Fan, Sakhila Ghimire, Misbah Tariq, Maria Drießlein, Marianna Maddaloni, Simon Holzinger, Gabriele Inchingolo, Lena Klostermeier, Claudia Gebhard, Nicholas Strieder, Daniela Hirsch, Daniela Weber, Elisabeth Meedt, Andreas Mamilos, Eva Vonbrunn, Maike Büttner-Herold, Ernst Holler, Andrea Ablasser, Matthias Edinger, Petra Hoffmann, Michael Rehli, Daniel Wolff, Wolfgang Herr, Sascha Göttert, Christian Schmidl, Hendrik Poeck

**Author notes:** These authors contributed equally (Chiara Suriano, Omer Khalid). Co-corresponding authors and shared senior authorship (Sascha Göttert, Christian Schmidl, Hendrik Poeck). Corresponding authors Sascha Göttert, Franz-Josef-Strauß-Allee 11, 93053 Regensburg, Telephone: 0941/944-38455. Christian Schmidl, Franz-Josef-Strauß-Allee 11, 93053 Regensburg. Hendrik Poeck, Franz-Josef-Strauß-Allee 11, 93053 Regensburg.

## Abstract

Allogeneic hematopoietic stem cell transplantation (allo-HSCT) is curative for hematological malignancies but limited by graft-versus-host disease (GvHD), in which donor T cells damage host tissues. Current prophylaxis broadly suppresses donor immunity, often compromising the beneficial graft-versus-leukemia (GvL) response, highlighting the need for strategies that uncouple GvHD from GvL. The cGAS–STING pathway can be strongly activated during conditioning-induced tissue damage and represents a potential therapeutic target. However, its context-dependent roles in inflammation and homeostasis have constrained its clinical translation. Here, we show that “timed” administration of the covalent STING inhibitor H151 before conditioning reduced GvHD-associated mortality without impairing GvL in murine allo-BMT models. Donor T cell activation and effector function were preserved, indicating that the protective effect operates at the level of GvHD target tissues rather than through systemic immunosuppression. Timed STING inhibition protected the intestinal epithelium by limiting apoptosis, preserving intestinal stem cell function, and sustaining metabolic fitness during conditioning-induced injury, independently of type I interferon (IFN-I) signaling. In allo-HSCT patients, low intestinal STING expression is associated with reduced transplant-related mortality. Together, these findings identify timed STING inhibition as a tissue-protective prophylactic strategy that could be incorporated into existing conditioning regimens to enhance efficacy while minimizing toxicity.

## Introduction

Allogeneic hematopoietic stem cell transplantation (allo-HSCT) is a curative therapy for hematological malignancies, with efficacy depending on the graft-versus-leukemia (GvL) effect mediated by donor T cells^1^. However, alloreactive donor T cells also drive graft-versus-host disease (GvHD), a major complication characterized by severe inflammation and tissue damage^2^. Acute GvHD predominantly targets the intestine, skin, and liver, leading to significant morbidity and mortality^3^. Intestinal GvHD causes extensive epithelial damage that disrupts the barrier and amplifies disease severity^4^. Current standard-of-care therapies broadly suppress donor T cell responses, often at the expense of GvL^5,6^. Achieving a clinically meaningful separation between GvHD and GvL therefore remains one of the central challenges in allo-HSCT.

The cyclic GMP–AMP synthase–stimulator of interferon genes (cGAS–STING) pathway senses cytosolic DNA released upon viral infection or cellular damage: cGAS generates cyclic GMP– AMP (cGAMP), which activates STING and triggers the production of type I interferon (IFN-I) and pro-inflammatory cytokines^7,8^. Beyond canonical signaling, STING also regulates cell death, autophagy, proliferation, and metabolic pathways, with distinct effects across biological settings^9–13^.

The functional outcomes of pathway activation are highly context dependent. In the intestine, STING and IFN-I contribute to epithelial homeostasis and barrier function^14,15^, while in tumor and immune cells STING activation promotes antitumor immunity^16–19^. In allo-HSCT, pharmacological STING agonism before transplantation alleviates intestinal GvHD via IFN-I signaling^20^, while STING expression in antigen-presenting cells constrains alloreactive T cell responses^21,22^. Conversely, sustained STING activation drives inflammatory and autoimmune disease^23–27^, prompting the development of pharmacological STING antagonists^28–31^. Conditioning regimens used in allo-HSCT induce extensive cellular damage and DNA release^32^. While this process can support antitumor responses^33^, it may also promote excessive STING activation, that amplifies inflammation and aggravates GvHD. We and others have shown that the pathway exerts divergent effects depending on the timing and context of its activation^20,22,34^. We therefore propose that transient inhibition to prevent hyperactivation in the conditioning phase may represent an exploitable therapeutic strategy in allo-HSCT.

Here, we show that administering the covalent STING inhibitor H151^35^ before conditioning mitigates GvHD without compromising GvL responses. The protective effect operates through preservation of intestinal epithelial integrity rather than direct suppression of donor T cells, involving reduced apoptosis and a preserved intestinal stem cell compartment independently of canonical IFN-I signaling. Sustained STING inhibition compromised this benefit. Together with patient data showing context-dependent correlations between intestinal STING expression and transplant-related mortality, these findings suggest temporally precise STING inhibition as a prophylactic strategy that protects tissues during conditioning while preserving alloreactive anti-tumor response.

## Methods

### Mice

BALB/c and C57BL/6J mice were purchased from Janvier-Labs. Lgr5-GFP reporter mice (B6.129P2-Lgr5tm1(cre/ERT2)Cle/J; RRID:IMSR_JAX:008875) were bred and maintained at the University Hospital Regensburg. Animals were housed under specific pathogen-free (SPF) conditions according to FELASA guidelines, with ad libitum access to food and water. Mice were acclimatized for at least one week before experimentation. All animal procedures were approved by the local regulatory authorities (Regierung von Oberbayern, ROB-55.2-2532.Vet_02-20-168). Experimental mice were 6–10 weeks old.

### H151 treatment

The covalent STING inhibitor H151 (InvivoGen, inh-h151) was used *in vivo* and *in vitro*. For *in vivo* administration, a 10 mg/mL stock solution in sterile DMSO was diluted in 10% Tween-80/PBS, and mice received 750 nmol H151 in 200 µL via intraperitoneal injection previously described^35^. Control animals received vehicle solution (10% Tween-80 in PBS). For *in vitro* assays (organoids and T cells), H151 was dissolved in DMSO and added to the culture medium at 0.5 µg/mL.

### Analysis of STING expression in allo-HSCT patients

GI biopsies were sampled from patients receiving allo-HSCT at different time points after transplantation and/or GvHD onset, and STING mRNA levels were measured by qPCR. 104 biopsies from 77 patients were analyzed. One biopsy per patient was selected for survival and outcome analysis: the biopsy obtained at GvHD onset, the first post-transplant biopsy in patients without GvHD or the only biopsy available. GI-aGvHD and clinical GvHD were graded according to the Lerner^36^ and system Glucksberg system^37^, respectively. Biopsies were classified as low (histological GvHD grade 0–2) or high (grade 3–4). The study was approved by the local ethics committee (University of Regensburg, 09/059 and 18-684482-101). All patients provided informed consent, and the study was conducted in accordance with the Declaration of Helsinki. Patient characteristics are listed in Table S6.

Remaining methods are listed in the Supplemental material.

## Results

### Timed pretransplant STING inhibition reduces experimental GvHD without impairing the GvL response

The conditioning regimens imply extensive genotoxic stress and tissue damage, which can aberrantly activate the cGAS–STING pathway and amplify GvHD^38,39^. To determine whether timed STING inhibition during this window could mitigate GvHD, we employed a major MHC-mismatched murine model of allogeneic bone marrow transplantation (allo-BMT) (C57BL/6 donors → BALB/c recipients). Mice received a single dose of the STING inhibitor H151 on the day of transplantation (day 0) three hours before conditioning **(Figure 1A)**. Timed STING inhibition reduced GvHD-associated weight loss **(Figure 1B)** and significantly improved overall survival **(Figure 1C)**.

**Figure 1.**
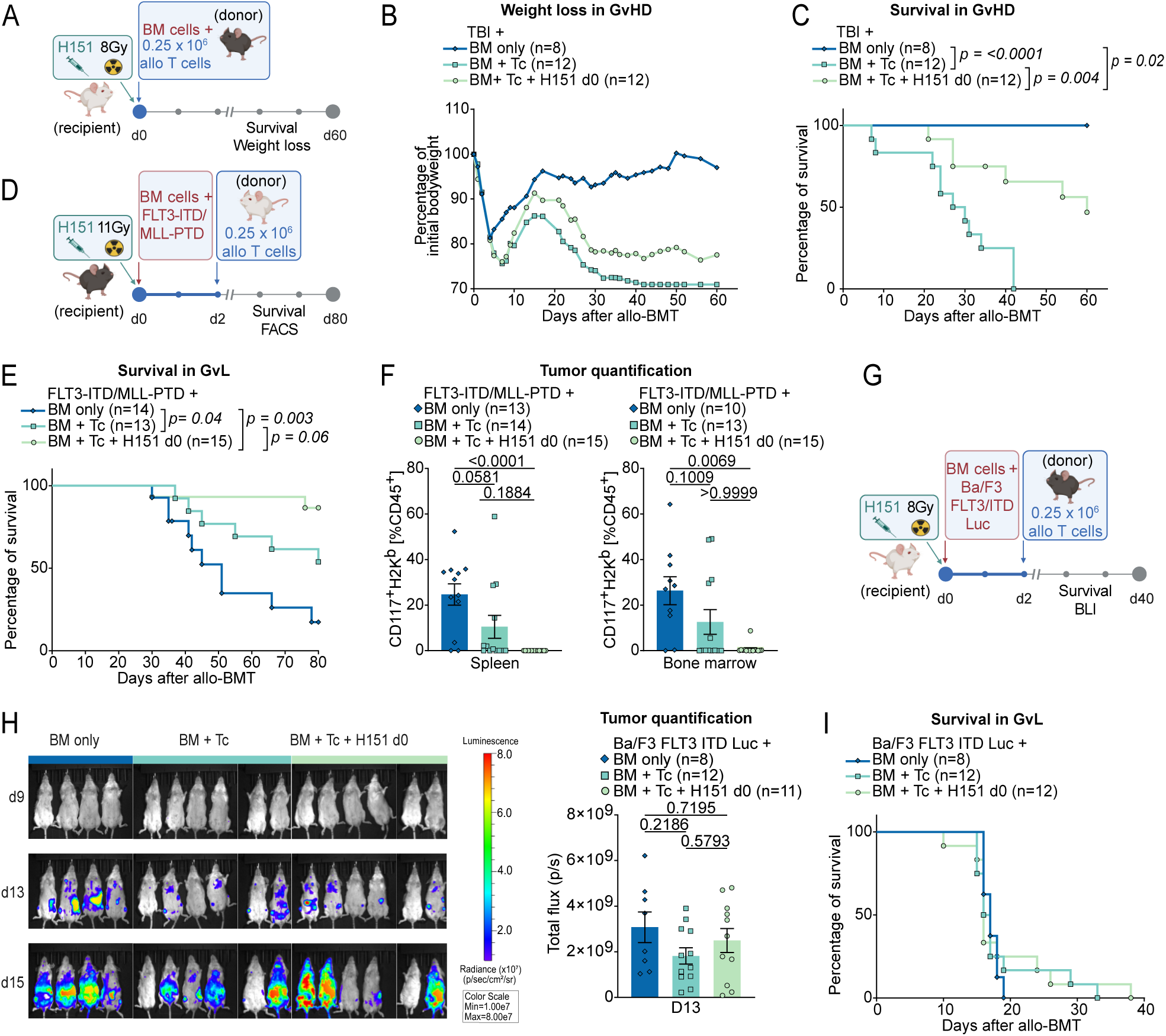
Timed pretransplant STING inhibition reduces experimental GvHD without impairing the GvL response. **(A)** Experimental design of the murine MHC major mismatch allo-BMT model: irradiated BALB/c recipient mice received allogeneic C57BL/6 bone marrow (BM) cells with or without additional C57BL/6 T cells (Tc), and with or without intraperitoneal (i.p.) administration of H151 3 hours prior to irradiation. **(B)** Body weight loss and **(C)** Kaplan-Meier survival analysis following allo-BMT. Data were pooled from two independent experiments. Survival was analyzed using the log-rank test. **(D)** Experimental design of the GvL-MLL model: irradiated C57BL/6 recipient mice received allogeneic BALB/c BM cells together with FLT3-ITD/MLL-PTD AML cells, with or without additional BALB/c Tc on day 2, and with or without i.p. H151 administration 3 hours before irradiation. **(E)** Survival of mice in the GvL model. Data was analysed by log-rank test from two independent experiments. **(F)** Leukemia cells (CD117+H-2Kb+) in spleen (left) and BM (right) were quantified by flow cytometry at human endpoint. Statistics by Kruskal-Wallis test with Dunn’s multiple comparisons test for nonparametric distribution. **(G)** Experimental design of the GvL-Ba/F3 model: irradiated BALB/c recipient mice received allogeneic C57BL/6 BM cells and Ba/F3 FLT3-ITD-luc⁺ leukemia cells, with or without additional C57BL/6 Tc on day 2, and with or without i.p. H151 administration 3 hours prior to irradiation. Leukemia progression was monitored by bioluminescence imaging (BLI). **(H)** Representative BLI images on days 9, 13, and 15, and quantification of BLI signal on day 13. Data were pooled from two independent experiments. P-values were calculated by one-way ANOVA with Tukey’s multiple comparisons test. **(I)** Kaplan– Meier survival analysis. Data were pooled from two independent experiments and were analyzed using the log-rank test. All data are shown as mean ± SEM. **(A) (D) (G)** were created in Biorender. **(A)** https://BioRender.com/0qxmk71 **(B)** https://BioRender.com/40irfe6 **(C)** https://BioRender.com/y9fdhl8

We next examined whether timed STING inhibition compromises the GvL effect. In a genetic acute myeloid leukemia (AML) model^40^, lethally irradiated recipients received allogeneic bone marrow with FLT3-ITD/MLL-PTD leukemia cells, with or without donor T cells **(Figure 1D).** As expected, donor T cells reduced leukemia burden compared to bone marrow–only controls, and this effect was maintained in H151-treated mice, as evidenced by a trend towards improved overall survival **(Figure 1E)**. Tumor burden in spleen and bone marrow at human endpoint **(Figure 1F**, gating strategy in **supplemental Figure 1A)** was then used to stratify lethality into GvHD or relapse: relapse was reduced by donor T cell infusion and further decreased in H151-treated mice **(supplemental Figure 1B)**, with unchanged GvHD severity across groups **(supplemental Figure 1C)**. We also tested the Ba/F3 FMS-like tyrosine kinase 3-internal tandem duplication-luciferase (Ba/F3 FLT3-ITD-Luc) model of acute lymphoid leukemia (ALL)^40^ **(Figure 1G)**. H151-treated mice showed comparable leukemic expansion by bioluminescence imaging **(Figure 1H; supplemental Figure 1D)** and survival **(Figure 1I)** to vehicle-treated controls.

Collectively, these data indicate that STING inhibition at the time of conditioning can uncouple GvHD from GvL, conferring protection from aGvHD without impairing the essential therapeutic anti-tumor immunity.

### Timed STING inhibition does not suppress T cell activation across pre-clinical and human systems

Donor T cells are central mediators of GvHD and key effectors of GvL. Given that timed STING inhibition reduced GvHD severity, we asked whether donor T cell function was altered.

To investigate the direct impact of STING inhibition on T cell responses, pan-T cells isolated from healthy donor PBMCs were stimulated with anti-CD3/CD28 beads and treated with H151 for 24 hours before single-cell RNA sequencing (scRNA-seq) **(Figure 2A**; sorting strategy in **supplemental Figure 2A)**. Graph-based clustering and cell type annotation identified major T cell subsets **(supplemental Figure 2B-C)**, and UMAP visualization showed overlapping distributions with no new clusters emerging in the H151 group **(Figure 2B)**. Gene set enrichment analysis (GSEA) reflected modest, subset-specific differences: metabolic pathways were enriched in effector-like populations while inflammatory, apoptotic, and T-cell activity programs were downregulated across multiple subsets. **(Figure 2C)**.

**Figure 2:**
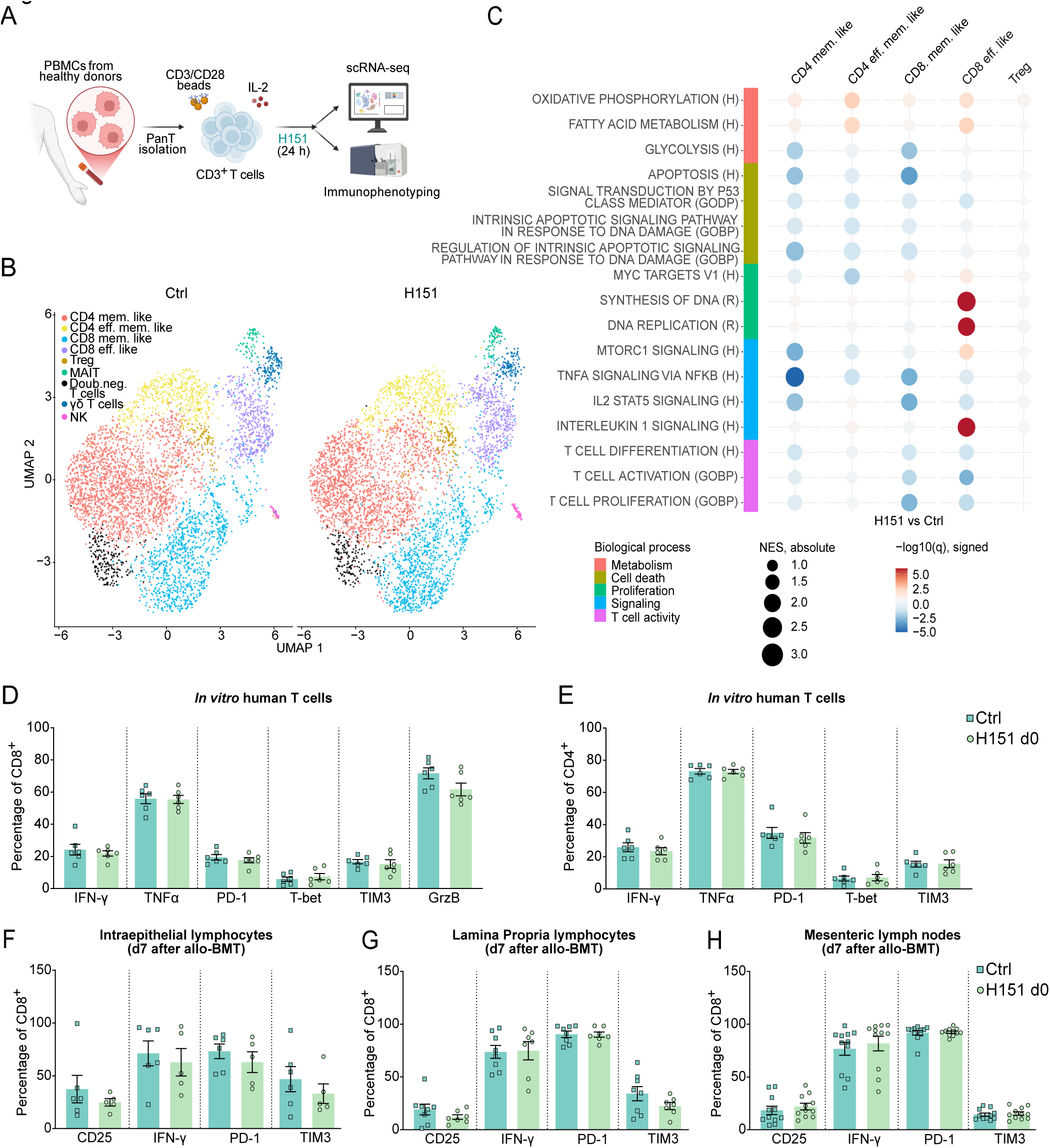
Timed STING inhibition does not suppress T cell activation across pre-clinical and human systems. **(A)** Experimental design. Human pan-T cells were isolated from healthy donors and stimulated with anti-CD3/CD28 beads, 30 U/mL human IL-2, and H151 for 24 h prior to scRNA-seq or flow cytometric analysis (n = 3, each group). Single cells for scRNA-seq were sorted as live/dead-CD45+ cells as indicated in **(supplemental Figure 2A)**. **(B)** UMAP plot of human pan-T scRNA-seq data colored by cell type annotation, split by experimental group. **(C)** Dot plot showing gene set enrichment analysis (GSEA) results for selected pathways across the indicated cell populations (H151 vs. control). Dot color represents the −log₁₀ of the GSEA q-value (FDR), with the sign indicating the direction of regulation (positive = upregulated, negative = downregulated). The scale is capped at -5/5 for better interpretability. Dot size reflects the GSEA normalized enrichment score (NES), indicating the magnitude of enrichment. Gene sets/pathways are derived from the Hallmark (H), Reactome (R), and GO Biological Process (GOBP) collections of MSigDB. Frequency of human CD8⁺ **(D)** and CD4⁺ **(E)** T cells positive for the indicated markers according to gates in **(supplemental Figure 2D)**. Data represents pooled samples from six healthy donors. Frequency of murine CD8⁺ T cells positive for the indicated markers, according to the gating strategy in **(supplemental Figure 3A)**. Cells were isolated from **(F)** intraepithelial lymphocytes (IEL), **(G)** lamina propria (LP), and **(H)** mesenteric lymph nodes (MLNs) of mice 7 days after allo-BMT. Data are pooled from three independent experiments. **(A)** was created with Biorender: https://BioRender.com/0whufjc

Functional validation by flow cytometry (gating strategy in **supplemental Figure 2D**) confirmed no remarkable alteration of T cell function, since H151 did not affect the frequency of CD8⁺ or CD4⁺ T cells producing cytokines or expressing activation markers **(Figure 2D-E; supplemental Figure 2E).**

To further evaluate the impact of STING inhibition on T cells *in vivo*, donor T cells were analyzed on day 7 after MHC-mismatched allo-BMT **(Figure 1A; supplemental Figure 3A)**. T cells were isolated from spleen, blood, mesenteric lymph nodes, and intestinal compartments (intraepithelial lymphocytes [IELs] and lamina propria [LP]). Frequencies of CD8+ (**Figure 2F-H; supplemental Figure 3B**) and CD4+ T cells **(supplemental Figure 3C)** expressing key activation and effector markers, and expression levels assessed by mean fluorescence intensity (MFI) **(supplemental Figure 3D-E)**, were comparable between groups across all tissues as well as in circulating and splenic T cells **(supplemental Figure 3D-E)**.

Together, these data argue against allogeneic T cells as the primary target of STING inhibition and indicate an alternative mechanism that protects against GvHD while retaining T cell activity. Despite preserved T cell activation, H151-treated mice showed reduced ileal T cell infiltration **(supplemental Figure 3F)**, leading our focus to the intestinal epithelium as potential site of the protective effect.

### Timed STING inhibition protects intestinal compartment from genotoxic stress through IFN-I–independent mechanisms

The intestinal epithelium is highly vulnerable to conditioning-induced injury, and its regenerative capacity is a critical determinant of barrier integrity and GvHD susceptibility^41–43^. Intestinal stem cells (ISCs) represent both the most sensitive population to genotoxic damage and the essential cellular source for barrier reconstruction. Given the established role of STING in epithelial homeostasis and regeneration, we hypothesized that this pathway may be particularly relevant in the regenerative ISC compartment.

In a human intestinal single-cell RNA-seq dataset^44^ cGAS and STING expression was higher in ISCs, transit-amplifying (TA) cells, and enterocyte progenitor compared to enterocytes, with higher expression of STING in the large intestine compared to small intestine **(supplemental Figure 4A–B, supplemental Table 4)**.

Given the high susceptibility of undifferentiated cells to genotoxic damage, we evaluated whether STING inhibition before conditioning could protect these populations. Mice were treated with H151 or vehicle before irradiation and intestinal crypts were isolated 24 hours later for scRNA-seq **(Figure 3A)**. Graph-based clustering, cell type annotation based on a reference dataset^45^, and established epithelial cell type markers, identified the major intestinal epithelial populations, with comparable distribution between conditions **(Figure 3B; supplemental Figure 4D-E)**. However, differential abundance analysis showed that one population (clusters 4 and 5 in **supplemental Figure 4D**) was less abundant in H151-treated mice **(supplemental Figure 4F)**. This population expressed higher levels of the intestinal stem cell markers *Lgr5* and *Olfm4* when compared to enterocyte-like cells (enterocyte progenitors and enterocytes), but lower levels of *Olfm4* compared to ISCs **(supplemental Table 4)**, with approximately 50% classified as stem cells by reference-based annotation with SingleR. Moreover, it also showed reduced enrichment of metabolic pathways alongside increased enrichment of DNA damage and chromatin remodeling gene sets **(supplemental Figure 4G)**, with elevated expression of general stress-response markers such as *Jun*, and markers of p53-driven stress responses such as *Mdm2* **(supplemental Figure 4H)**. We therefore annotated this population as stressed progenitors, consistent with previous reports describing the response of the intestinal epithelium to irradiation damage^46–48^. The reduced abundance of stressed progenitors in H151-treated mice indicates that STING inhibition limits the proportion of cells undergoing acute stress responses after irradiation.

**Figure 3.**
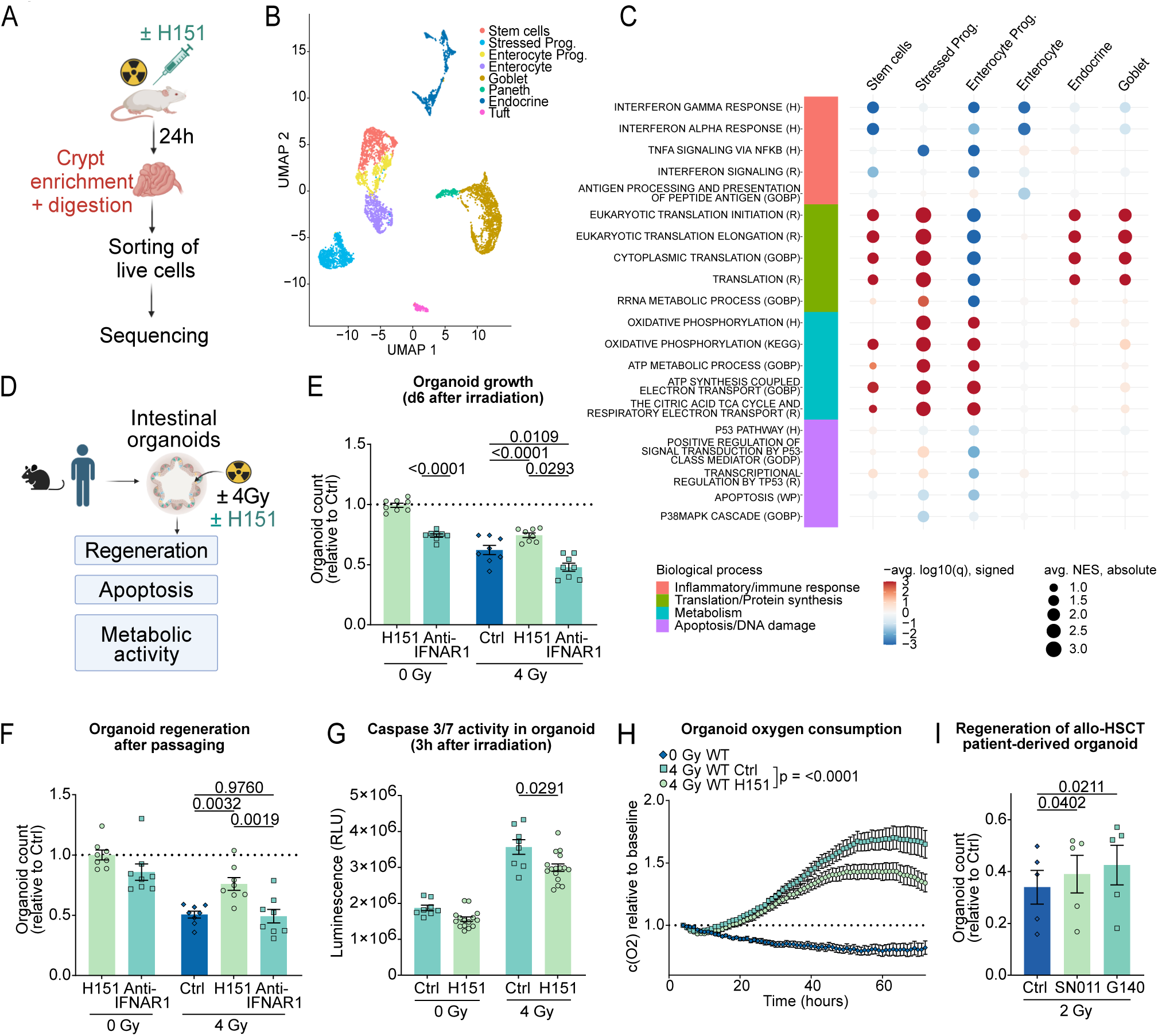
Timed STING inhibition protects intestinal compartment from genotoxic stress through IFN-I–independent mechanisms. **(A)** Experimental design for single-cell RNA sequencing (scRNA-seq) of intestinal crypt cells isolated from irradiated BALB/c mice treated with H151 or vehicle in a single experiment with n = 3 mice per group. UMAP plot of murine intestinal scRNA-seq data colored by cell type annotation, from all experimental conditions **(B)** and split by experimental group **(supplemental Figure 4E)**. **(C)** Dot plot showing gene set enrichment analysis (GSEA) results for selected pathways across the indicated cell populations (H151 vs. control). Dot color represents the negative of the average of the log_10_ GSEA q-values (FDR) across the random sampling runs (see supplemental methods), with the sign indicating the direction of regulation (positive = upregulated, negative = downregulated). The scale is capped at -3/3 for better interpretability. Dot size reflects the average normalized enrichment score (NES) across random runs, indicating the magnitude of gene set enrichment. Gene sets/pathways are derived from the Hallmark (H), Reactome (R), GO Biological Process (GOBP), and KEGG collections of MSigDB. **(D)** Experimental overview of organoid assays. Organoid counts were assessed prior to passaging **(E)** and after passaging **(F)**, under steady-state conditions or following 4 Gy irradiation, with or without H151 or IFNAR1 antibody. Data were normalized to control (dotted line) and pooled from four independent experiments (n=8 mice). Statistical analysis was performed using one sample t-test **(E,** left) or oneway ANOVA with Tukey’s multiple comparisons test **(E,** right; **F)**. **(G)** Quantification of apoptotic activity in organoids using the Caspase-Glo 3/7 assay under steady-state conditions or following 4 Gy irradiation, with or without H151. Statistical analysis was performed using unpaired t-test with Welch’s correction. **(H)** Measurement of metabolic activity by oxygen concentration in media of H151-treated or control organoids alone or following irradiation with 4 Gy. Data were pooled from 3 independent experiments (n=8 wells). P-values were calculated by unpaired t-test with Welch’s correction between AUC (area under curve). **(I)** Growth of human large intestine organoids derived from allo-HSCT patients, treated with the STING inhibitor SN011 or the cGAS inhibitor G140, with or without 2 Gy irradiation. Data were pooled from five donors across two independent experiments. Statistical analysis was performed using repeated-measures one-way ANOVA with uncorrected Fisher’s LSD test. All data are presented as mean ± SEM. **(A) (D)** were created in Biorender: **(A)** https://BioRender.com/ky7t242 **(D)** https://BioRender.com/170lrp3

GSEA comparing H151-treated and control mice across epithelial populations revealed additional pathway-level differences **(Figure 3C)**: in ISCs and enterocyte-like cells, H151 was associated with depletion of inflammatory/immune response gene sets, particularly those related to interferons. Pathways associated with translation capacity were enriched in most cell types except for enterocyte-like cells, while aerobic metabolic pathways were upregulated across all undifferentiated populations. Apoptotic signaling appeared to be downregulated in enterocyte progenitors specifically. Together, these observations suggest a less inflammatory and more regenerative epithelium state in H151-treated mice.

To functionally test these observations, we used murine intestinal organoid cultures, which replicate epithelial self-renewal and regenerative properties *ex vivo*, assessing viability at day 6 and regenerative capacity after passaging **(Figure 3D)**. Under steady-state conditions, a single H151 dose did not impair organoid viability, but following irradiation, H151 preserved organoid numbers **(Figure 3E)**, and this protective effect was more pronounced upon passaging **(Figure 3F)**. To also investigate whether these effects involved the canonical STING–IFN-I signaling axis, we included treatment with an IFNAR-1 blocking antibody. IFNAR-1 blockade reduced organoid viability under steady-state conditions, consistent with the established role of IFN-I in epithelial integrity^20^. In contrast to H151, IFNAR-1 blockade did not prevent acute toxicity and worsened organoid regeneration following irradiation **(Figure 3E-F)**, indicating that intact IFN-I signaling is also required for recovery from genotoxic stress. The opposite phenotype of H151 demonstrates that its protective action is not mediated by suppression of downstream IFN-I signaling but through STING-specific, non-canonical mechanisms. The reduced IFN-I response detected by scRNA-seq in H151-treated mice **(Figure 3C)** is therefore likely a consequence rather than the driver of epithelial protection. Consistently, the cGAS inhibitor RU521 partially recapitulated the regenerative benefit **(supplemental Figure 4I-J)**. Moreover, H151 protected organoids from the clinically relevant chemotherapeutics cisplatin and busulfan **(supplemental Figure 4K-L)**, extending the protective effect to additional conditioning modalities.

We next investigated potential non-canonical mechanisms: Caspase 3/7 measurement showed that H151 significantly reduced apoptosis in irradiated organoids **(Figure 3G),** and continuous oxygen monitoring revealed that H151-treated organoids maintained significantly higher metabolic activity than irradiated controls **(Figure 3H)**.

Finally, to assess whether this protective mechanism is conserved in human intestinal tissue, we examined the effect of STING inhibition in large-intestinal organoids derived from allo-HSCT patients. Remarkably, H151 before irradiation showed no protective effect **(supplemental Figure 4M)**, in line with reported cell type-specific differences in H151 efficacy^49^. However, the alternative STING inhibitor SN011^50^ and cGAS inhibitor G140^51^ both improved organoid regeneration following irradiation **(Figure 3I)**. Although the effect was modest, these findings suggest that the protective mechanism is conserved in the human intestine and highlights its potential translational relevance.

In summary, timed STING inhibition promotes epithelial survival and regeneration under genotoxic stress through IFN-I-independent mechanisms, providing a mechanistic rationale for improved transplant outcomes.

### Timed STING inhibition supports intestinal integrity during conditioning and allo-BMT

Having shown that timed STING inhibition protects intestinal epithelial cells *in vitro*, we validated these findings *in vivo*. In a syngeneic BMT model, H151 or vehicle was administered before irradiation **(Figure 4A)**, and small intestinal crypts isolated seven days later were transferred to organoid cultures. Crypts from H151-treated mice generated significantly more organoids than controls **(Figure 4B)**, indicating direct mitigation of radiation-induced epithelial damage.

**Figure 4.**
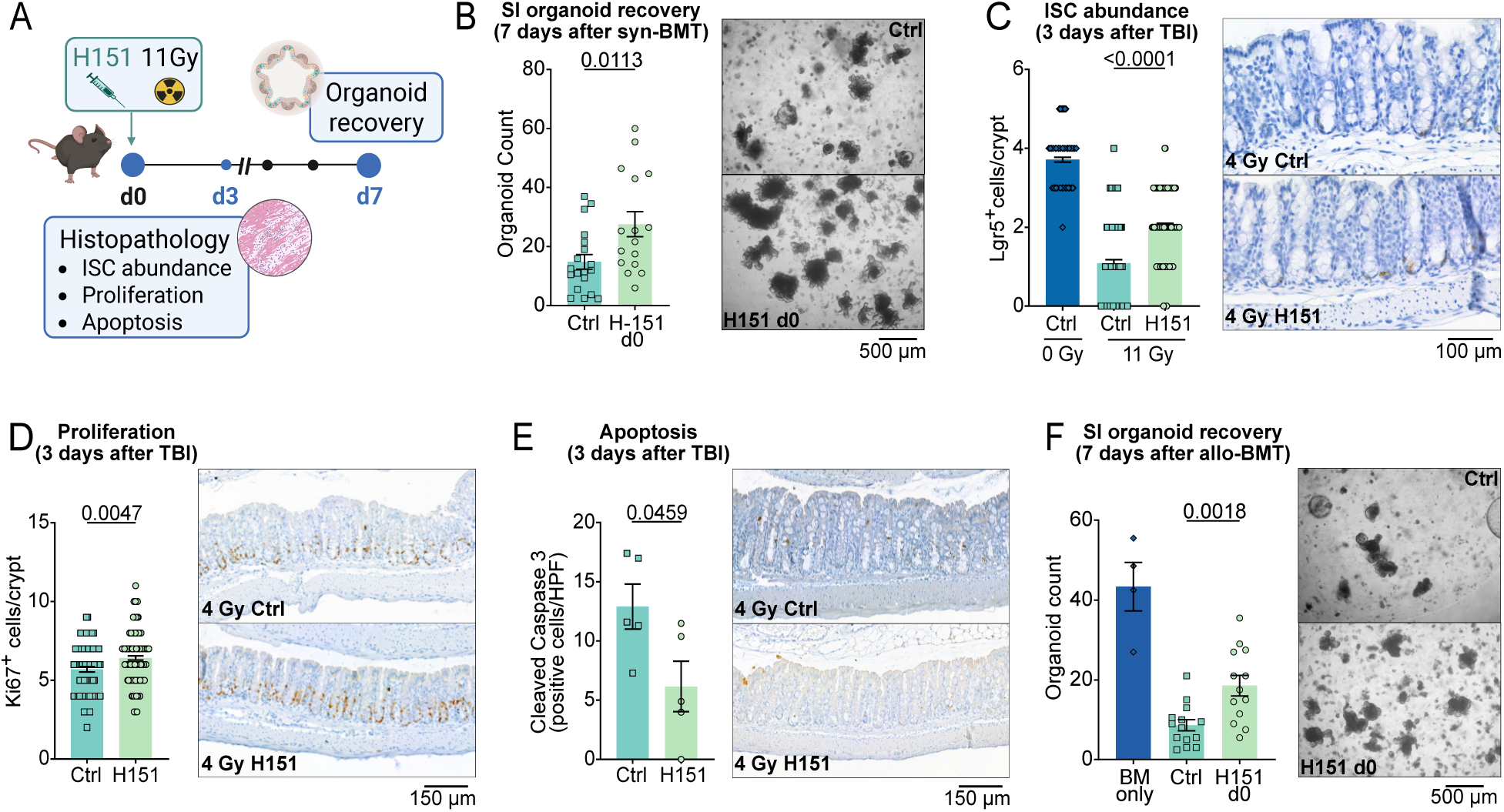
Timed STING inhibition supports intestinal integrity during conditioning and allo-BMT. **(A)** Experimental design. **(B)** Number of recovered small intestinal organoids on day 7 after total body irradiation (TBI) followed by transplantation of syngeneic BM cells, with representative images. Data were pooled from 5 independent experiments (n = 18 Ctrl; n = 15 H151 d0). Statistical analysis by Mann–Whitney test for nonparametric data. Histopathological score and representative images of **(C)** Lgr5+ cells, **(D)** Ki67+ cells and **(E)** cleaved caspase 3-positive cells in the colon on day 3 after TBI in mice treated with H151 or vehicle. Data were pooled from two independent experiments and crypts were analysed in 5 mice per group. The number of crypts was in **(C)** n = 139 (Ctrl 0 Gy), n = 161 (Ctrl 4 Gy) and 116 (H151 d0); in **(D)** n = 84 (Ctrl) and 144 (H151 d0). In **(E)** n = 5 mice (Ctrl) and 5 mice (H151 d0). Scale bars: 100 µm in **(C)** and 150 µm in **(D–E)**. Statistical analysis: Mann–Whitney test for nonparametric data **(C–D)**; unpaired t-test with Welch’s correction **(E)**. **(F)** Number of recovered small intestinal organoids on day 7 after allo-BMT, with representative images (scale bar: 500 µm). Data were pooled from 4 independent experiments (n = 4 BM only; n = 14 Ctrl; n = 13 H151 d0). Statistical analysis by one-way ANOVA with Tukey’s multiple comparisons test, followed by unpaired t-test comparing Ctrl vs H151 d0. **(A)** was created with Biorender: https://BioRender.com/522zv8z

To gain further mechanistic insights, mice were sacrificed for histological analysis three days after total body irradiation (TBI) with or without H151 pre-treatment. TBI markedly depleted Lgr5+ ISCs in vehicle-treated controls, while H151 partially preserved Lgr5+ abundance **(Figure 4C)**. H151-treated mice also displayed increased Ki67+ proliferation **(Figure 4D)** and reduced cleaved caspase-3 staining **(Figure 4E)**, recapitulating the *in vitro* observations.

Finally, we tested whether this protective effect extends to allo-BMT, where conditioning-induced epithelial damage is compounded by alloreactive T cell–mediated inflammation. BALB/c recipients were conditioned and transplanted with allogeneic BM alone or together with donor T cells, with H151 administered before irradiation. Donor T cells markedly impaired crypt-derived organoid recovery seven days later; however, H151 pre-treatment significantly restored organoid formation **(Figure 4F)**.

Collectively, these data validate the epithelial-protective effect of timed STING inhibition *in vivo* and demonstrate that a single pre-conditioning dose preserves ISC function, limits epithelial apoptosis, and supports regeneration following both conditioning alone and allo-BMT.

### Intestinal STING expression in allo-HSCT patients correlates with TRM

We next asked whether intestinal STING expression is associated with clinical outcomes. STING mRNA levels were determined in gastrointestinal (GI) biopsies from 77 allo-HSCT patients (patient characteristics in **supplemental Table 5**). High STING expression was associated with increased TRM (transplant-related mortality) risk **(Figure 5A)**. A deeper stratification by histological GvHD grade (low: Lerner 0-2; high: Lerner 3-4) revealed that in patients with low-grade GvHD, high STING expression was significantly associated with worse TRM **(Figure 5B)**, supporting STING inhibition as a prophylactic strategy. In contrast, in the small subset with high-grade GvHD, the association was reversed, with higher STING expression correlating with reduced TRM **(Figure 5C)**. Although this must be interpreted cautiously given the limited sample size, it raises the hypothesis that STING may adopt a context-specific role under conditions of severe inflammation, potentially contributing to tissue repair or the restoration of immune homeostasis during hyperinflammatory states.

**Figure 5.**
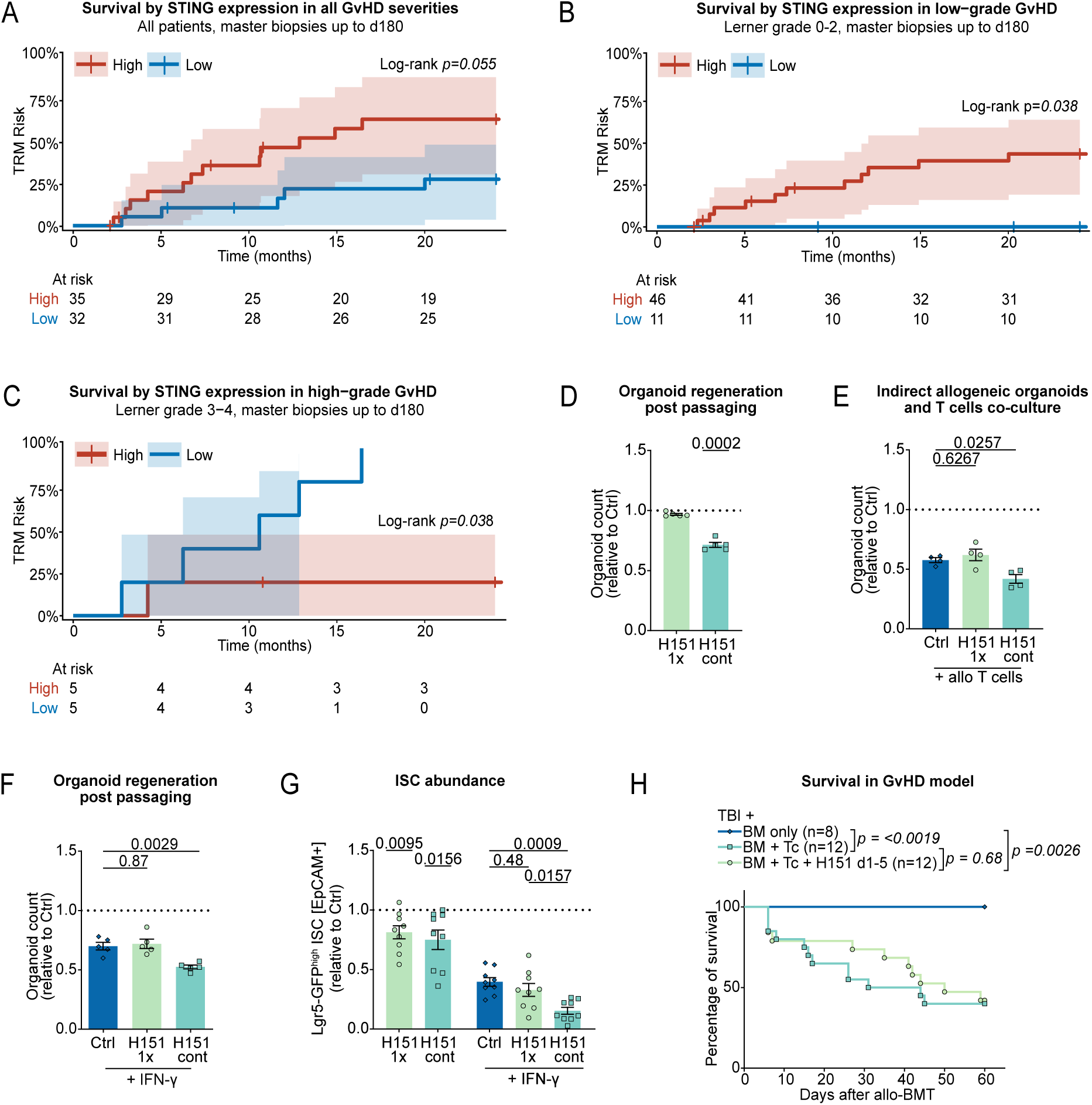
STING exerts a time- and context-dependent role in epithelial regeneration and survival of patients after allo-HSCT. **(A-B-C)** Kaplan–Meier overall survival of allo-HSCT patients stratified by STING expression in large intestine biopsies collected up to day 180 post-transplant, with corresponding numbers at risk. **(A)** Entire cohort; **(B)** patients with low-grade GvHD defined by histological Lerner grade 0-2; **(C)** patients with high-grade GvHD defined by histological Lerner grade 3-4. **(D)** Murine small intestinal (SI) organoid counts after passaging under steady-state conditions following single (1X) or repeated H151 treatment (days 0, 3, and 5 post-seeding). Data were normalized to control (dotted line) and pooled from two independent experiments. Statistical analysis by one-sample t-test. **(E)** Murine SI organoids co-cultured with activated splenic T cells in surrounding media (indirect co-culture with no direct contact) for 4 days. Organoid growth was assessed as in **(D).** Data were pooled from four independent experiments and p-values were calculated by ordinary one-way ANOVA with Dunnett’s multiple comparisons test. **(F)** SI organoids were challenged with IFN-γ on days 3 and 5 post-seeding and treated with single (d0) or repeated H151 (days 0, 3, and 5) or vehicle. Regenerative capacity was assessed after passaging by organoid counts. Data were normalized to control (dotted line), pooled from two independent experiments, and analyzed by one-way ANOVA with Dunnett’s multiple comparisons test. **(G)** Intestinal stem cell (ISC) frequency assessed by flow cytometry (gating strategy shown in **supplemental Figure 5A**) in Lgr5-GFP organoids under steady-state conditions or IFN-γ stimulation, with single or repeated H151 treatment as in **(F)**. Data from seven independent experiments are shown. Statistical analysis by one sample t-test **(G,** left) and one-way ANOVA with Tukey’s multiple comparisons test **(G,** right). **(H)** Kaplan–Meier survival analysis in the murine allo-BMT model of mice receiving H151 i.p. from day 1 to day 5. Recipient mice here received double dose of allogeneic T cells (0.5 x 10^6^) compared to the model in (Figure 1A). Data were pooled from three independent experiments and were analyzed using the log-rank test. All data are shown as mean ± SEM.

### STING inhibition has a time- and context-dependent effect on epithelial regeneration and fails to improve outcomes when applied after allo-BMT

The context-dependent pattern across inflammatory states in patients prompted us to assess whether sustained pharmacological STING inhibition may be detrimental under heightened inflammation. To model this, organoids were cultured under steady-state conditions or with allogeneic T cells or IFN-γ, applying single or repeated H151 treatment. Repeated treatment under steady-state conditions caused more pronounced impairment of regeneration than single-dose treatment, suggesting that prolonged STING blockade compromises baseline regenerative capacity **(Figure 5D)**. When mimicking the inflammatory environment of allo-HSCT with allogeneic T cells in the surrounding media or with direct IFN-γ exposure, continuous H151 further worsened organoid regeneration **(Figure 5E–F)**. Given that IFN-γ is a major mediator of intestinal stem cell injury during alloreactive inflammation^52^, we directly examined Lgr5-GFP^high^ EpCAM⁺ ISC abundance in Lgr5-GFP organoids (gating in **supplemental Figure 5A**). The ISC pool was markedly depleted by IFN-γ exposure; single-dose H151 did not rescue ISC abundance while continuous treatment further worsened this effect **(Figure 5G)**. This contrasted with the preservation of Lgr5⁺ ISCs after single-dose H151 *in vivo* **(Figure 4C)**, reflecting the different nature of stem cell injury in each setting: while STING inhibition limits radiation-induced injury, IFN-γ depletes ISCs through mechanisms that are not reversed by timed STING inhibition. Together, these findings indicate that sustained STING inhibition sensitizes epithelium to inflammatory stress, in line with the reversed STING-TRM association in high-grade GvHD patients where STING activity appears to exert a protective rather than pathogenic effect.

Notably, the enhanced *in vitro* sensitivity to sustained STING inhibition was not recapitulated *in vivo*. Daily H151 administration from day +1 to day +5 after allo-BMT failed to improve survival **(Figure 5H)** but did not worsen it, demonstrating that low post-transplant STING activity early after allo-BMT is tolerated. Together, these findings indicate that the protective effect of STING inhibition critically depends on timing and inflammatory context. Transient inhibition during the conditioning window protects the intestinal epithelium from acute genotoxic injury, while sustained blockade compromises ISC resilience and loses the survival benefit. The clinical observation that intestinal STING expression correlates with transplant-related mortality in a context-dependent manner further supports a model in which the pathway can exert either pathogenic or protective functions depending on the context of activation.

## Discussion

Innate immune sensors are well-established drivers of GvHD initiation: conditioning-induced cellular damage releases DAMPs and PAMPs that activate these sensors, fueling cytokine production and donor T cell priming^38,53,54^. The cGAS–STING pathway is therefore a strong candidate for sensing cytosolic DNA released following conditioning and initiating inflammatory cascades^55^. In allo-HSCT, previous efforts to exploit this pathway have focused on STING agonists given before transplantation, which induce protective IFN-I responses. More broadly, both STING agonists and antagonists have entered clinical evaluation, demonstrating the feasibility and safety of pharmacological STING targeting in humans^56–58^. Agonists are being developed to enhance antitumor and antiviral immunity^59–63^, while antagonists have shown benefits in inflammatory and autoimmune diseases, where chronic STING inhibition attenuates inflammation without compromising immune competence^35,64–67^. Our approach instead exploits brief STING inhibition during the early phase of conditioning-induced injury, recapitulating these anti-inflammatory properties through direct effects on the intestinal epithelium. H151 prevented apoptosis of intestinal stem cells, consistent with reports that STING activation can drive apoptosis, necroptosis, and other cell death programs under genotoxic stress^9,23,68^. Single-cell transcriptomic analysis of the irradiated epithelium showed that H151 depleted inflammatory and immune response pathways in ISCs and progenitor cells, while modestly attenuating pro-apoptotic programs, primarily in enterocyte progenitors. Concurrently, aerobic metabolic pathways were enriched, supporting a shift toward a regeneration-competent state. By limiting these pro-apoptotic and inflammatory programs, timed STING inhibition may reduce DAMP/PAMP release and direct antigen presentation by intestinal epithelial cells, the latter shown to contribute to GvHD initiation through MHC class II–dependent mechanisms^69^. This places tissue protection upstream of the T cell response: rather than dampening alloreactivity, STING inhibition reduces the inflammatory and antigenic signals that fuel it, providing a mechanistic basis for the observed uncoupling of GvHD from GvL.

A frequent concern about modulating innate immunity in allo-HSCT is that reducing GvHD will inevitably compromise GvL, as most current prophylactic strategies dampen donor T cell responses^70,71^. Our human T cell experiments excluded broad suppression of activation by STING inhibition, with scRNA-seq revealing only modest, subset-specific differences. This is consistent with recent work in CAR-T cell therapy, showing that T cell–intrinsic genetic STING ablation can foster responses against solid tumors when combined with a STING agonist, but does not impact baseline T cell function^72^. Transient pharmacological inhibition, such as ours, would therefore not be expected to cause major T cell changes. Indeed, donor T cells from H151-treated mice retained normal activation despite the suppressed intestinal inflammation observed early after conditioning. Beyond direct T cell effects, timed STING inhibition may also influence T cells via antigen-presenting cell (APC) function: genetic STING deficiency has been shown to prolong APC survival and enhances donor T cell expansion^21,22^. Although our intervention is temporary, it may transiently mimic this effect, providing a potential explanation for the moderately improved GvL response in our FLT3-ITD/MLL-PTD model.

Our data also show that the protective effect of STING inhibition is not mediated by IFN-I, distinguishing our approach from STING agonism. The latter relies on canonical IFN-I signaling, which supports intestinal epithelial homeostasis and barrier integrity^14,15^ and has been shown to mitigate GvHD when applied before transplantation^20^. Despite this mechanistic difference, both approaches share a key feature: a single pre-conditioning dose confers durable benefit. This common outcome is explained by the temporal dynamics of STING activation during conditioning. Within hours of irradiation, DNA damage and cytosolic DNA release drive a wave of STING activation that contributes to injury of the radiosensitive ISC compartment. STING agonism applied before transplantation pre-loads protective IFN-I signaling so that it is already engaged when this damage occurs, whereas pre-conditioning H151 blocks the acute STING activation, preventing the initial wave of epithelial apoptosis and subsequent GvHD priming. Once this critical phase has passed, STING can resume its homeostatic and regenerative functions -a role independently characterized in a parallel study from our group, which demonstrated that STING is required for intestinal stem cell–driven regeneration triggered by a microbial metabolite, independently of IFN-I signaling^73^. Of note, although sustained STING inhibition sensitizes ISCs to IFN-γ–mediated damage *in vitro*, daily H151 administration outside the conditioning window (day 1 to 5) was not detrimental *in vivo* but also did not provide therapeutic benefit. The consequences of STING modulation are therefore not intrinsic to the pathway itself but shaped by the timing of intervention and the ongoing inflammatory state.

The clinical relevance of this context-dependent biology is reflected in our patient data. In the overall cohort, low STING expression was associated with reduced TRM, supporting STING inhibition in allo-HSCT. However, stratification by GvHD severity refined this interpretation: in patients with low-grade GvHD and limited intestinal inflammation, low STING expression conferred a survival benefit, in line with the efficacy of prophylactic but not post-transplant STING inhibition in our model. In contrast, in the highly inflammatory setting of high-grade GvHD, STING appeared required. Together, this temporal and context-dependent pattern support a prophylactic strategy focused on preserving epithelial barrier integrity rather than immunosuppression and indicates limited utility once GvHD is established. The concept of prophylactic strategies preserving epithelial integrity before transplantation has shown clinical benefit in other settings employing keratinocyte growth factor^74^ or regulatory T cell-based approaches^75,76^, both of which promote tissue protection and modulate alloreactivity. Our approach complements these efforts without requiring graft manipulation and integrating directly with existing conditioning regimens. Moreover, post-transplant interventions promoting GvL through STING activation, such as Menin inhibition^77^, further reinforce restricting STING inhibition to the prophylactic window. The rapid plasma clearance of H151 ^35^ would naturally confine its effects to the peri-conditioning period, thereby preserving STING-dependent homeostatic and regenerative functions during later transplant stages.

In summary, our study demonstrates that timed STING inhibition uncouples GvHD from GvL by preventing conditioning-induced epithelial injury through non-canonical, IFN-I-independent mechanisms. These findings refine our understanding of STING biology in allo-HSCT, identify timing as a central determinant of pathway modulation outcomes, and propose pharmacological strategies enabling precise temporal control as a path toward improved patient outcomes. This precision modulation could be integrated with existing into existing conditioning and GvHD prophylaxis regimens to enhance efficacy while minimizing toxicity. Future studies will need to address optimal dosing, the long-term impact on immune reconstitution, and potential synergies with established conditioning and GvHD prophylaxis regimens including post-transplant cyclophosphamide (PTCy) or other immunosuppressants.

## Acknowledgement

This study was supported by the Deutsche Forschungsgemeinschaft (DFG, German Research Foundation) projektnummer 324392634-TRR 221 (to H.P., C.Sc., E.H., M.B-H., M.E., D.We., P.Ho., M.R., D.Wo., W.H.); Projektnummer 395357507 – SFB 1371 (to H.P., E.H., E.M.), the Wilhelm Sander Foundation (2021.040.1 to H.P), the Else-Kröner-Fresenius-Stiftung (funding line: Else-Kröner Forschungskolleg to E.M.), The Bavarian State Ministry of Science and Art (COVID funds to H.P.). H.P. is supported by the EMBO Young Investigator Program, by the European Union (project MICROBOTS, Grant No. 101124680 to H.P.). Views and opinions expressed are however those of the author(s) only and do not necessarily reflect those of the European Union or the European Research Council. Neither the European Union nor the granting authority can be held responsible for them. This work was performed in the framework of the National Cancer Therapy Center (NCT) WERA. This work is part of the doctoral thesis of C.Su. and O.K. at the University of Regensburg.

Special thanks to Maria Krieger (Poeck group) and Doris Gaag (Pathology UKR) for the precious technical support. Single cell RNA sequencing experiments were conducted at the NGS Unit of the Leibniz Institute of Immunotherapy (LIT) of Regensburg, Germany. We thank Hanna Stanewsky and Johanna Raithel for their technical assistance. FACS and sorting were performed at the FACS core of the LIT (Regensburg, Germany) with the technical help of Irina Fink, Jaqueline Dirmeier and Niklas Wenzl. Schematics were generated with Biorender.

## Authorship

Conceptualization: S.G., C.Sc., H.P. Data curation: C.Su., O.K., P.H., S.G., C.Sc., H.P. Formal Analysis: C.Su., O.K., P.H., S.H., S.G., C.Sc., H.P. Investigation: C. Su., O.K., K.F., S.Gh, M.T., M.D., M.M., G.I., L.K., D.H., A.M., E.V., S.G. Methodology: P.H., S.H., C.G., N.S., D.H., D.We., E.M., M.B-H., E.H., A.A., M.E., P.Ho., M.R., D.Wo. Project administration: S.G., C.Sc., H.P. Resources: M.B-H., E.H., A.A., M.E., P.Ho., M.R., D.Wo., W.H., C.Sc., H.P. Supervision: S.G., C.Sc., H.P. Visualization: C.Su., O.K., P.H., S.H., S.G., C.Sc., H.P. Writing – original draft: C.Su., O.K., S.G. Writing – editing: S.G., C.Sc., H.P. Writing – review: all authors. Funding acquisition: C.Sc., H.P. Contributions are specified according to CRediT (Contributor Roles Taxonomy). All authors reviewed and approved the final version of the manuscript.

## Data Sharing Statement

For original data, please contact the lead contact

## Bibliography

1. Giralt S, Bishop MR. Principles and overview of Allogeneic Hematopoietic Stem Cell Transplantation. Cancer Treat Res. 2009;144:1–21.

2. Negrin RS. Graft-versus-Host Disease versus Graft-versus-Leukemia. Hematology Am Soc Hematol Educ Program. 2015;2015:225–30.

3. Zeiser R, Blazar BR. Acute Graft-versus-Host Disease — Biologic Process, Prevention, and Therapy. N Engl J Med. 2017;377(22):2167–2179.

4. Takatsuka H, Iwasaki T, Okamoto T, Kakishita E. Intestinal graft-versus-host disease: mechanisms and management. Drugs. 2003;63(1):1–15.

5. Martin PJ, Rizzo JD, Wingard JR, et al. First- and Second-Line Systemic Treatment of Acute Graft-versus-Host Disease: Recommendations of the American Society of Blood and Marrow Transplantation. Biol Blood Marrow Transplant. 2012;18(8):1150–1163.

6. Malard F, Holler E, Sandmaier BM, Huang H, Mohty M. Acute graft-versus-host disease. Nat Rev Dis Primers. 2023;9(1):27.

7. Diner EJ, Burdette DL, Wilson SC, et al. The Innate Immune DNA Sensor cGAS Produces a Noncanonical Cyclic Dinucleotide that Activates Human STING. Cell Rep. 2013;3(5):1355–1361.

8. Dong M, Fitzgerald KA. DNA-sensing pathways in health, autoinflammatory and autoimmune diseases. Nat Immunol. 2024;25(11):2001–2014.

9. Hayman TJ, Baro M, MacNeil T, et al. STING enhances cell death through regulation of reactive oxygen species and DNA damage. Nat Commun. 2021;12(1):2327.

10. Vila IK, Chamma H, Steer A, et al. STING orchestrates the crosstalk between polyunsaturated fatty acid metabolism and inflammatory responses. Cell Metab. 2022;34(1):125–139.e8.

11. Liu D, Wu H, Wang C, et al. STING directly activates autophagy to tune the innate immune response. Cell Death Differ. 2019;26(9):1735–1749.

12. Ranoa DRE, Widau RC, Mallon S, et al. STING Promotes Homeostasis via Regulation of Cell Proliferation and Chromosomal Stability. Cancer Res. 2019;79(7):1465–1479.

13. Vila IK, Guha S, Kalucka J, Olagnier D, Nadine Laguette. Alternative pathways driven by STING: From innate immunity to lipid metabolism. Cytokine Growth Factor Rev. 2022;68:54–68.

14. Katlinskaya Y V, Katlinski K V, Lasri A, et al. Type I Interferons Control Proliferation and Function of the Intestinal Epithelium. Mol Cell Biol. 2016;36(7):1124–1135.

15. Canesso MCC, Lemos L, Neves TC, et al. The cytosolic sensor STING is required for intestinal homeostasis and control of inflammation. Mucosal Immunol. 2018;11(3):820–834.

16. Woo SR, Fuertes MB, Corrales L, et al. STING-Dependent Cytosolic DNA Sensing Mediates Innate Immune Recognition of Immunogenic Tumors. Immunity. 2014;41(5):830–842.

17. Marcus A, Mao AJ, Lensink-Vasan M, Wang L, Vance RE, Raulet DH. Tumor-Derived cGAMP Triggers a STING-Mediated Interferon Response in Non-tumor Cells to Activate the NK Cell Response. Immunity. 2018;49(4):754–763.e4.

18. Li W, Lu L, Lu J, et al. cGAS-STING–mediated DNA sensing maintains CD8+ T cell stemness and promotes antitumor T cell therapy. Sci Transl Med. 2020;12(549):eaay9013.

19. Linder A, Nixdorf D, Kuhl N, et al. STING activation improves T-cell-engaging immunotherapy for acute myeloid leukemia. Blood. 2025;145(19):2149–2160.

20. Fischer JC, Bscheider M, Eisenkolb G, et al. RIG-I/MAVS and STING signaling promote gut integrity during irradiation- and immune-mediated tissue injury. Sci Transl Med. 2017;9(386):eaag2513.

21. Bader CS, Barreras H, Lightbourn CO, et al. STING differentially regulates experimental GVHD mediated by CD8 versus CD4 T cell subsets. Sci Transl Med. 2020;12(552):eaay5006.

22. Wu Y, Tang CA, Mealer C, et al. STING negatively regulates allogeneic T cell responses by constraining antigen-presenting cell function. Cell Mol Immunol. 2021;18(3):632–643.

23. Ding Z, Wang R, Li Y, Wang X. MLKL activates the cGAS-STING pathway by releasing mitochondrial DNA upon necroptosis induction. Mol Cell. 2025;85(13):2610–2625.e5.

24. Konno H, Chinn IK, Hong D, et al. Pro-inflammation Associated with a Gain-of-Function Mutation (R284S) in the Innate Immune Sensor STING. Cell Rep. 2018;23(4):1112–1123.

25. Jeremiah N, Neven B, Gentili M, et al. Inherited STING-activating mutation underlies a familial inflammatory syndrome with lupus-like manifestations. J Clin Invest. 2014;124(12):5516–5520.

26. Liu Y, Jesus A A, Marrero B, et al. Activated STING in a Vascular and Pulmonary Syndrome. N Engl J Med. 2014;371(6):507–518.

27. Pan Y, You Y, Sun L, et al. The STING antagonist H-151 ameliorates psoriasis via suppression of STING/NF-κB-mediated inflammation. Br J Pharmacol. 2021;178(24):4907–4922.

28. Prabakaran T, Troldborg A, Kumpunya S, et al. A STING antagonist modulating the interaction with STIM1 blocks ER-to-Golgi trafficking and inhibits lupus pathology. EBioMedicine. 2021;66:103314.

29. Guo F, Zhang J, Gao Y, et al. Discovery and Total Synthesis of Anhydrotuberosin as a STING Antagonist for Treating Autoimmune Diseases. Angew Chem Int Ed. 2025;64(1):e202407641.

30. Zhou P, Yang G, Wang Y, et al. Development of indole derivatives as inhibitors targeting STING-dependent inflammation. Bioorg Med Chem. 2025;126:118216.

31. Zhou X, Yue H, Wang X, et al. Discovery of 3,4-dihydroisoquinoline-2(1H)-carboxamide STING inhibitors as anti-inflammatory agents. Eur J Med Chem. 2025;297:117922.

32. Ramadan A, Paczesny S. Various Forms of Tissue Damage and Danger Signals Following Hematopoietic Stem-Cell Transplantation. Front Immunol. 2015;Volume 6- 2015.

33. Deng L, Liang H, Xu M, et al. STING-Dependent Cytosolic DNA Sensing Promotes Radiation-Induced Type I Interferon-Dependent Antitumor Immunity in Immunogenic Tumors. Immunity. 2014;41(5):843–852.

34. Fischer JC, Bscheider M, Göttert S, et al. Type I interferon signaling before hematopoietic stem cell transplantation lowers donor T cell activation via reduced allogenicity of recipient cells. Sci Rep. 2019;9(1):14955.

35. Haag SM, Gulen MF, Reymond L, et al. Targeting STING with covalent small-molecule inhibitors. Nature. 2018;559(7713):269–273.

36. Lerner KG, Kao GF, Storb R, Buckner CD, Clift RA, Thomas ED. Histopathology of graft-vs.-host reaction (GvHR) in human recipients of marrow from HL-A-matched sibling donors. Transplant Proc. 1974;6(4):367–71.

37. Glucksberg H, Storb R, Fefer A, et al. Clinical manifestations of graft-versus-host disease in human recipients of marrow from HL-A-matched sibling donors. Transplantation. 1974;18(4):295–304.

38. Härtlova A, Erttmann SF, Raffi FA, et al. DNA Damage Primes the Type I Interferon System via the Cytosolic DNA Sensor STING to Promote Anti-Microbial Innate Immunity. Immunity. 2015;42(2):332–343.

39. Hill GR, Crawford JM, Cooke KR, Brinson YS, Pan L, Ferrara JL. Total Body Irradiation and Acute Graft-Versus-Host Disease: The Role of Gastrointestinal Damage and Inflammatory Cytokines. Blood. 1997;90(8):3204–3213.

40. Rückert T, Andrieux G, Boerries M, et al. Human β-defensin 2 ameliorates acute GVHD by limiting ileal neutrophil infiltration and restraining T cell receptor signaling. Sci Transl Med. 2022;14(676):eabp9675.

41. Kim CK, Yang VW, Bialkowska AB. The Role of Intestinal Stem Cells in Epithelial Regeneration Following Radiation-Induced Gut Injury. Curr Stem Cell Rep. 2017;3(4):320–332.

42. Hill GR, Ferrara JL. The primacy of the gastrointestinal tract as a target organ of acute graft-versus-host disease: rationale for the use of cytokine shields in allogeneic bone marrow transplantation. Blood. 2000;95(9):2754–2759.

43. Johansson JE, Brune M, Ekman T. The gut mucosa barrier is preserved during allogeneic, haemopoietic stem cell transplantation with reduced intensity conditioning. Bone Marrow Transplant. 2001;28(8):737–742.

44. Wang Y, Song W, Wang J, et al. Single-cell transcriptome analysis reveals differential nutrient absorption functions in human intestine. J Exp Med. 2020;217(2):e20191130.

45. Haber AL, Biton M, Rogel N, et al. A single-cell survey of the small intestinal epithelium. Nature. 2017;551(7680):333–339.

46. Murata K, Jadhav U, Madha S, et al. Ascl2-Dependent Cell Dedifferentiation Drives Regeneration of Ablated Intestinal Stem Cells. Cell Stem Cell. 2020;26(3):377–390.e6.

47. Ayyaz A, Kumar S, Sangiorgi B, et al. Single-cell transcriptomes of the regenerating intestine reveal a revival stem cell. Nature. 2019;569(7754):121–125.

48. Yui S, Azzolin L, Maimets M, et al. YAP/TAZ-Dependent Reprogramming of Colonic Epithelium Links ECM Remodeling to Tissue Regeneration. Cell Stem Cell. 2018;22(1):35–49.e7.

49. Chan R, Cao X, Ergun SL, et al. Cysteine allostery and autoinhibition govern human STING oligomer functionality. Nat Chem Biol. 2025;21(10):1611–1620.

50. Hong Z, Mei J, Li C, et al. STING inhibitors target the cyclic dinucleotide binding pocket. Proc Natl Acad Sci U S A. 2021;118(24):e2105465118.

51. Lama L, Adura C, Xie W, et al. Development of human cGAS-specific small-molecule inhibitors for repression of dsDNA-triggered interferon expression. Nat Commun. 2019;10(1):2261.

52. Takashima S, Martin ML, Jansen SA, et al. T cell–derived interferon-γ programs stem cell death in immune-mediated intestinal damage. Sci Immunol. 2019;4(42):eaay8556.

53. Penack O, Holler E, van den Brink MR. Graft-versus-host disease: regulation by microbe-associated molecules and innate immune receptors. Blood. 2010;115(10):1865–1872.

54. Heidegger S, van den Brink MR, Haas T, Poeck H. The Role of Pattern-Recognition Receptors in Graft-Versus-Host Disease and Graft-Versus-Leukemia after Allogeneic Stem Cell Transplantation. Front Immunol. 2014;Volume 5-2014.

55. Brennan T V, Rendell VR, Yang Y. Innate Immune Activation by Tissue Injury and Cell Death in the Setting of Hematopoietic Stem Cell Transplantation. Front Immunol. 2015;Volume 6-2015.

56. Luke JJ, Pinato DJ, Juric D, et al. Phase I dose-escalation and pharmacodynamic study of STING agonist E7766 in advanced solid tumors. J Immunother Cancer. 2025;13(2):e010511.

57. Jiang M, Chen P, Wang L, et al. cGAS-STING, an important pathway in cancer immunotherapy. J Hematol Oncol. 2020;13(1):81.

58. Le Naour J, Zitvogel L, Galluzzi L, Vacchelli E, Kroemer G. Trial watch: STING agonists in cancer therapy. Oncoimmunology. 2020;9(1):1777624.

59. Gallovic MD, Junkins RD, Sandor AM, et al. STING agonist-containing microparticles improve seasonal influenza vaccine efficacy and durability in ferrets over standard adjuvant. J Control Release. 2022;347:356–368.

60. Humphries F, Shmuel-Galia L, Jiang Z, et al. A diamidobenzimidazole STING agonist protects against SARS-CoV-2 infection. Sci Immunol. 2021;6(59):eabi9002.

61. Fu J, Kanne DB, Leong M, et al. STING agonist formulated cancer vaccines can cure established tumors resistant to PD-1 blockade. Sci Transl Med. 2015;7(283):283ra52–283ra52.

62. Pan BS, Perera SA, Piesvaux JA, et al. An orally available non-nucleotide STING agonist with antitumor activity. Science. 2020;369(6506):eaba6098.

63. Gehrcken L, Deben C, Smits E, Van Audenaerde JRM. STING Agonists and How to Reach Their Full Potential in Cancer Immunotherapy. Adv Sci (Weinh). 2025;12(17):e2500296.

64. Domizio J Di, Gulen MF, Saidoune F, et al. The cGAS–STING pathway drives type I IFN immunopathology in COVID-19. Nature. 2022;603(7899):145–151.

65. Gulen MF, Samson N, Keller A, et al. cGAS–STING drives ageing-related inflammation and neurodegeneration. Nature. 2023;620(7973):374–380.

66. Alee I, Chantawichitwong P, Leelahavanichkul A, Paludan SR, Pisitkun T, Pisitkun P. The STING inhibitor (ISD-017) reduces glomerulonephritis in 129.B6.Fcgr2b-deficient mice. Sci Rep. 2024;14(1):11020.

67. Han X, Ma G, Peng R, et al. Discovery of an Orally Bioavailable STING Inhibitor with In Vivo Anti-Inflammatory Activity in Mice with STING-Mediated Inflammation. J Med Chem. 2025;68(3):2963–2980.

68. Sun Y, Aliyari SR, Parvatiyar K, et al. STING directly interacts with PAR to promote apoptosis upon acute ionizing radiation-mediated DNA damage. Cell Death Differ. 2025;32(6):1167–1179.

69. Koyama M, Mukhopadhyay P, Schuster IS, et al. MHC Class II Antigen Presentation by the Intestinal Epithelium Initiates Graft-versus-Host Disease and Is Influenced by the Microbiota. Immunity. 2019;51(5):885–898.e7.

70. Hadjis AD, Nunes NS, Khan SM, et al. Post-Transplantation Cyclophosphamide Uniquely Restrains Alloreactive CD4+ T-Cell Proliferation and Differentiation After Murine MHC-Haploidentical Hematopoietic Cell Transplantation. Front Immunol. 2022;Volume 13- 2022.

71. Makuuchi Y, Nakashima Y, Nishimoto M, Koh H, Hino M, Nakamae H. Posttransplant cyclophosphamide contributes to the impairment of the graft-versus-leukemia effect and the amelioration of graft-versus-host disease with the suppression of alloreactive T cells in a murine stem cell transplant model. Exp Hematol. 2023;123:56–65.

72. Piseddu I, Endres R, Lanzl F, et al. STING Ablation in T Cells Is Required for the Efficacy of STING Agonists in CAR-T Cell Immunotherapy of Pancreatic Cancer. Gastroenterology. Published online May 10, 2026.

73. Göttert S, Thiele Orberg E, Fan K, et al. The microbial metabolite desaminotyrosine protects against graft-versus-host disease via mTORC1 and STING-dependent intestinal regeneration. Nat Commun. 2025;16(1):9282.

74. Schulz E, Curtis LM, Holtzman NG, et al. Phase 1/2 study of high-dose palifermin for GVHD prophylaxis in patients undergoing HLA-matched unrelated donor HCT. Blood. 2025;146(8):944–950.

75. Edinger M, Hoffmann P, Ermann J, et al. CD4+CD25+ regulatory T cells preserve graft-versus-tumor activity while inhibiting graft-versus-host disease after bone marrow transplantation. Nat Med. 2003;9(9):1144–1150.

76. Meyer EH, Salhotra A, Gandhi AP, et al. Orca-T vs allogeneic hematopoietic stem cell transplantation (Precision-T): a multicenter, randomized phase 3 trial. Blood. 2026;147(11):1168–1177.

77. Fetsch V, Schwöbel L, Ozyerli-Goknar E, et al. Menin inhibition enhances graft-versus-leukemia effects by T-cell activation and endogenous retrovirus induction in AML. Blood. 2026;147(5):584–601.

