## Supplemental material for "Timed STING Inhibition Mitigates Gastrointestinal GvHD While Preserving Graft-versus-Leukemia Activity After Allo-HSCT"

### Methods

#### Murine Graft-versus-host disease model

To induce GvHD after allo-BMT, we employed a major mismatch model (H-2kd/H-2kb) with myeloablative conditioning. BALB/c recipients were irradiated with 8 Gy TBI and 3-4 hours later, transplanted intravenously with $0.5x10^6$ allogeneic (C57BL/6J) T-cell-depleted bone marrow cells, either alone or in combination with $0.25x10^6$ purified allogeneic T cells. Bone marrow cells were isolated from naïve C57BL/6J mice. Donor $CD4^+$ and $CD8^+$ T cells were isolated from pooled spleens of naïve C57BL/6 mice using magnetic bead separation (Miltenyi Biotec) and mixed at a 1:1 ratio. Recipient mice were monitored daily in a non-blinded manner and euthanized when reaching predefined human endpoint criteria: 1) more than 30% weight loss; 2) severe diarrhea with bleeding; 3) apathy or other criteria indicated by standard institutional guidelines.

#### Murine Graft-versus-leukemia model

Two previously described GvL models were used[1] . In one, BALB/c mice received 8 Gy TBI 750 followed by transplantation of $0.5 \times 10^6$ C57BL/6 BM cells and 500- Ba/F3 leukemia cells (DSMZ ACC 300) transfected with FLT3-ITD and luciferase (luc)+. Two days after allo-BMT, $0.25 \times 10^6$ donor T cells were injected. BM-only control mice did not receive T cells. Leukemia progression was monitored by *in vivo* bioluminescence imaging (BLI).
In the second model, C57BL/6 recipients were transplanted with BALB/c BM cells together with 5000-7500 FLT3-ITD/MLL-PTD AML cells, with or without additional $0.25 \times 10^6$ BALB/c T cells on day 2. Leukemic burden ($CD117^+$H-2K^$b^+$ cells) in BM and spleen was assessed by flow cytometry. Leukemia cells were kindly provided by Natalie Köhler.

#### Bioluminescence imaging (BLI)

For *in vivo* BLI, mice received intraperitoneal (i.p.) injections of luciferin (150 mg/kg body weight) and were anesthetized by inhalation with 2.5% isoflurane. Imaging was performed with the IVIS 50 bioluminescence imaging system (Xenogen) and quantified using the Living image Software (Xenogen).

#### *In vitro* T cell activation

Human pan-T cells were isolated from PBMCs of healthy donors using the Pan T Cell Isolation Kit (Miltenyi Biotec). Cells ($1 \times 10^5$/well) were cultured in 96-well U-bottom plates in RPMI 1640 medium supplemented with 10% FCS, 2 mM L-glutamine, 100 U/mL penicillin, and 30 U/mL recombinant human IL-2 (PeproTech). For stimulation, cells were incubated with anti-CD3/CD28 Dynabeads™ (Thermo Fisher) in the presence or absence of H151 (0.5 µg/mL) for 24 h at 37 °C and 5% $CO_2$. After incubation, cells were collected for flow cytometry analysis or for single-cell RNA sequencing (scRNA-seq).

#### Flow cytometry

For intracellular cytokine analysis, cells were stimulated for 3 h with Cell Stimulation Cocktail containing protein transport inhibitors (eBioscience, 00-4975-93). Surface and intracellular staining were performed using fluorochrome-conjugated antibodies listed in supplemental Table 1. Data were acquired on BD LSR Fortessa X-20 or Symphony A5 cytometers and analyzed with FlowJo software (TreeStar). All protocols included live/dead staining and, for murine samples, CD16/CD32 Fc blocking. Surface staining was performed for 30 min at 4 °C. For intracellular staining, cells were fixed and permeabilized with the Foxp3/Transcription Factor Staining Buffer Set (eBioscience) and stained for 40 min at 4 °C.

##### Analysis of published human intestine scRNA-seq data

For the assessment of STING and cGAS gene expression in human intestinal cell types, a published scRNA-seq dataset from the human intestine has been used (GEO accession number GSE125970)[2]. Briefly, for each intestinal segment (ileum, colon, rectum), cells of two human donors have been processed on a 10x Genomics Chromium device to obtain transcriptomic profiles, resulting in 14537 cells after QC filtering. For the differential gene expression analysis of STING and cGAS between cell types of a given intestinal segment, and for the comparison of cells from different intestinal segments, the Wilcox test (Seurat 5.0.1) has been applied to the library-size normalized and log(n+1) transformed data. P values have been corrected for multiple testing with the FDR approach, per intestinal segment/across all comparisons between intestinal segments. Cell type annotations from the published data have been used for this analysis. Only data for ileum and colon are shown.

##### Crypt enrichment for single-cell RNA sequencing

The small intestine of irradiated mice was isolated 16 hours after irradiation. It was then flushed with PBS, opened longitudinally, content carefully removed and cut into 1cm pieces. The pieces were washed with cold PBS until the supernatant was clear and then incubated for 35 minutes at 4°C in PBS + 5mM EDTA. Intestinal crypts were isolated in fractions in PBS. Therefore, crypts were forcefully shaken twice and the supernatant passed through one 100µm and two 70µm strainers. This procedure was repeated four times. Crypt enriched fractions were combined and digested with TrypleE Express Enzyme (Gibco) containing 0.1% BSA for 5 minutes at 37°C. The reaction was stopped by adding PBS and cells passed through a 40µm strainer. Single cell suspensions were stained and sorted as indicated above.

##### Cell sorting for single-cell RNA sequencing

Human PanT cells were treated as described above. Afterwards, single cell suspensions were stained with CD45-APC (Biolegend) and respective hashing antibody (TotalSeq™-C0251 anti-human Hashtag 1 and TotalSeq™-C0253 anti-human Hashtag 4 Antibody, Biolegend). Live/dead staining was performed immediately before sorting by adding propidium iodide. Single cells were sorted as live/dead-CD45+ cells, as shown in supplemental Figure 2A.

For the intestinal epithelial cells, single cell suspensions were stained with 0.5 mg/ml propidium iodide and sorted for live cells, as shown in supplemental Figure 4C.

#### Single-cell RNA sequencing

10x processing and sequencing

For the generation of scRNA-seq libraries from murine intestinal cells, cells were processed on a 10x Chromium Controller using the On-Chip Multiplexing technology with the Chromium GEM-X OCM 3' Chip Kit v4 4-plex (10x Genomics, Cat. No. 1000747) and the GEM-X Universal 3' Gene Expression v4 4-plex Kit (Cat. No. 1000779), following the manufacturer's protocol. For each mouse sample, a total of 20,000 cells (determined by FACS counting) were loaded onto the chip and distributed across two lanes with 10,000 cells per lane. Each chip comprised four lanes and included one control mouse sample and one H151-treated mouse sample to mitigate batch effects. Amplified cDNA was used to generate 3′ scRNA-seq libraries, which were quantified using a Qubit™ Fluorometer (ThermoFisher) and quality-checked on a TapeStation (Agilent). Libraries were sequenced on an Illumina NovaseqX Plus at Novogene GmbH (Munich, Germany) using PE sequencing (150 cycles each direction).

To produce scRNA-seq libraries from samples of in-vitro activated human T cells, individual treatment-related samples from each donor were labeled with hashtag-oligo (HTO) antibodies (see above). Subsequently, they were loaded onto a Chromium Next GEM Chip K using the Chromium Next GEM Single Cell 5′ Reagent Kit v2 and the Chromium Single Cell 5′ Feature Barcode Kit (Dual Index) (10x Genomics) according to the manufacturer's protocol. Amplified cDNA was used for both 5′ gene expression library generation using the Chromium Single Cell 5′ Library Kit v2 and cell surface protein (CSP) library preparation using the Chromium Single Cell 5′ Feature Barcode Kit (10x Genomics). This resulted in one scRNA and one CSP library per donor. 5′ scRNA and CSP libraries were quantified using a Qubit™ Fluorometer (Thermo Fisher Scientific) and quality-checked using a TapeStation system (Agilent Technologies). Libraries were pooled at a ratio of 1 (CSP): 3 (scRNA) and sequenced on an Illumina NextSeq 2000 platform on a P3 flow cell, and with 100% scRNA on a second P3 flow cell. All sequencing runs were performed with a PE-26-10-10-92 sequencing setup. The individual samples are distinguished during analysis on the single-cell level via the HTO barcodes.

Raw read processing

For the scRNA-seq of murine intestinal cells, demultiplexing was performed at Novogene GmbH (Munich, Germany) with the bcl2fastq2 software (v2.20). Using the multi command from Cellranger (version 9.0.1) with standard options, sequencing reads were then mapped to the mouse genome (GRCm39), cells were called, and UMI count files for the gene expression libraries were created. The multi config csv file for each library was set up to map the different on-chip-multiplexing barcoding groups.

For the scRNA-seq of *in-vitro* samples of human T cells, demultiplexing of sequencing reads was carried out using bcl2convert provided by Illumina Dragen (v-3.8.4, NextSeq 2000, Illumina Inc., San Diego, California, USA). Sequencing reads were combined and mapped to the human genome (GRCh38) using the Cellranger (version 8.0.1) count command with standard options and expected cells per library set to 36000. Count files for gene expression and HTO libraries were created by Cellranger.

#### Data analysis of single cell RNA-seq

Further processing and analysis of single cell RNA-seq data was performed mainly in R (version 4.3.0), and primarily with the Seurat package (version 5.0.1). The same processing has been applied to both the murine intestine data and the in-vitro human T cell data, if not explicitly stated otherwise.

Preprocessing

For the murine intestinal data, doublets were identified using the scDblFinder R-package[3] (version 1.14.0, used together with SingleCellExperiment, version 1.22.0) and excluded from the analysis. scDblFinder was run in random mode individually on the cell barcodes resulting from each lane of the GEM-X 3' OCM chips used. Cells have additionally been filtered to have a total UMI count and number of features above 1000 and 200, respectively. The maximum percentage of mitochondrial genes was set to 5%. This resulted in 41095 cells remaining for analysis (59285 cells initially called by Cellranger).

For the human T cell data, to assign treatment groups to single cells based on HTO counts, the HTODemux() function from the Seurat package was employed with default settings (positive-quantile parameter of 0.99) after centralized log normalization. Cells that could not be assigned to a treatment group due to insufficient HTO labeling (negative cells) were excluded from the analysis, as well as HTO-identified doublets (cells with ambiguous labeling). Additionally, cells from groups unrelated to this study were also excluded based on their HTO label. Further filtering has been performed for a total UMI count and number of features above 1000 and 200, respectively, and a maximum percentage of mitochondrial genes of 5%. In total, this resulted in 28624 remaining cells for analysis.

Clustering and annotation

UMI counts were normalized for library size and log(n+1) transformed. After selecting the 2000 top variable features, data was then mean-centered, scaled to unit variance and subjected to principal component analysis (PCA). For the human T cell in-vitro data, cell cycle effects were additionally regressed out using UCell[4] scores (S-phase/G2M signatures from Tirosh *et al.*[5]) prior to PCA. Subsequently, the principal components (PCs) were used as input to the harmony R-package[6] (version 1.2.0) to integrate data from the different libraries (default parameters).

Afterwards, graph-based clustering was performed on a shared nearest-neighbor graph constructed from the first 20 dimensions of the harmony-transformed data. Clustering (standard Louvain algorithm) was run at different resolutions to find the optimal clustering setup. For visualization, a UMAP (Uniform Manifold Approximation and Projection[7]) dimensionality reduction was additionally applied to the first 20 dimensions of the harmony-transformed data.

For the human T cell in-vitro data, a clustering resolution of 1.2 was chosen to be able to separate the relevant T cell subtypes. Annotation was then done using a combination of automatic annotation with STCAT[8] (version 1.0.8, Python 3.9.16, R 4.4.3, Seurat 5.3.0), and assessment of standard marker gene expression.

For the murine intestinal data, a clustering resolution of 0.3 was initially chosen. Cells were annotated broadly into epithelial cells, T cells and monocytes/B cells via assessment of marker gene expression and automatic annotation with SingleR[9] (version 2.2.0). SingleR was used together with a scRNA-seq reference dataset from the murine intestine (GEO accession number GSE92332, full length atlas data[10]) to annotate epithelial cell types. Only the epithelial subset of cells was used for subsequent analysis. Variable feature selection (2000), scaling, PCA and harmony integration were repeated, and clustering and UMAP were computed again on the first

20 dimensions of the harmony embedding. A clustering resolution of 0.5 was chosen for detailed annotation of epithelial cells via marker genes and SingleR annotation. Additionally, for annotating the cell population identified as Stressed Progenitors, stress and DNA damage response marker genes as well as differential gene expression and gene set enrichment analysis results (see respective sections for details) from comparison with the other epithelial cell populations were used for annotation. A small cluster of epithelial-like cells that could not be properly categorized and partly also showed T cell marker expression was excluded from the analysis.

Differential gene expression analysis

Within annotated cell populations, testing for differential gene expression between treatment groups (Control and H151) was performed on the raw UMI count data with the NEBULA R-package[11] (version 1.5.3). NEBULA fits (by default) a negative binomial mixed model to count data and estimates both cell- and subject-level overdispersions. For each cell population, a NEBULA negative binomial gamma mixed model was fitted to the data. Library size was set as the scaling factor (normalization), and minimum counts per cell for a gene to be tested were set to 0.005. The fit was computed with the NEBULA-LN method. All other parameters were set to default values. For the murine intestinal data, explanatory variables were treatment group, scRNA-seq library and FACS processing batch. Each mouse was defined as a subject (mixed model random effect). For the human in-vitro T cell data, treatment group was set as the explanatory variable of the model, while donor (equals library) was defined as subject. For some of the annotated cell populations, there were major differences in the numbers of cells across the different replicates. Therefore, for differential expression analysis between treatment groups, replicates with higher cell numbers were downsampled to avoid biases resulting from replicate domination. Downsampling was performed without replacement, to at maximum two (human in-vitro T cell data) or three times (murine intestine data) the number of cells of the replicate with the lowest cell number.

When testing for differential gene expression between annotated cell populations, irrespective of treatment group, the Wilcoxon Rank Sum test from Seurat's FindMarkers() function was used on the log-normalized data, employing FDR correction for multiple testing. To be as unrestrictive as possible, all genes expressed in at least 1% of cells in either of the two compared cell populations were included in testing.

Gene set enrichment analysis

For pathway/gene set analysis, the GSEA (gene set enrichment analysis)[12] implementation from the fgsea R-package (version 1.26.0) was employed. As input for GSEA, the negative $\log_{10}$ p-values signed with the direction of the log fold-change (FC) were used. If there were p-values with a value of zero, these were set to the smallest non-zero normalized floating-point number ($2.23 \cdot 10^{-308}$) prior to log-transformation. Values were then sorted by the signed log-transformed p-values in descending order, breaking possible ties by sorting according to log FC. Gene sets were obtained from MSigDB (Molecular signatures database)[13] via the msigdbr R-package (version 7.5.1). Chosen gene set collections were Hallmark gene sets (H), KEGG, Wikipathways and Reactome subsets from curated gene sets (C2), and GO Biological Process from GO gene sets (C5). Only gene sets where at least 70% of genes were in the genes tested for differential expression in each comparison were assessed. The p-values obtained from GSEA for the different comparisons and gene sets were corrected for multiple testing with the false discovery rate (FDR) approach, controlling for an FDR of 10%. For comparisons between treatment groups within a given cell population, correction was performed across all hypotheses within a given comparison.

For comparisons between cell populations, correction was performed across all hypotheses associated with a given comparison of cell populations.

Repeated random downsampling for differential expression and GSEA analysis

For the murine intestine data, the down sampling process for differential expression analysis between treatment groups yielded only small cell numbers for some of the annotated populations. Thus, this process was repeated ten times with different random draws to obtain stable results. For each random draw of cells, differential expression analysis and GSEA between treatment groups has been performed as described above. Analysis was focused on gene sets that were reliably up/- or downregulated at an FDR of 10% in at least 70% of the random runs in a given cell population and comparison of treatment groups. For reporting GSEA results from repeated random sampling, the average of the log(FDR) and the NES (normalized enrichment score) across random runs were computed.

Differential abundance analysis

Analysis for differential cell type abundance (DA) between treatment groups was performed with the MASS R-package (version 7.3.60)[14]. Briefly, for annotated cell populations from the murine intestine scRNA-seq data, a negative binomial model with the variable's treatment group, library and FACS processing batch was fitted to the cell count data. Total cell count per sample (i.e., library size) was used as an offset for normalization. P-values were derived by performing a Wald test with the respective linear contrasts of model parameter estimates. They were then adjusted for multiple testing using the FDR approach.

#### Induction of intestinal damage by total body irradiation or allo-BMT

For irradiation-induced damage, BALB/c mice received 8 Gy total body irradiation (TBI) with the Gammacell 40 exactor, while C57BL/6 mice received 11 Gy. Where indicated, mice were subsequently injected intravenously with $5 \times 10^6$ syngeneic (C57BL/6) bone marrow (BM) cells.

For induction of allogeneic tissue damage, Balb/c recipient mice were subjected to 8 Gy TBI followed by intravenous injection of $5 \times 10^6$ allogeneic (BALB/c) BM cells, either alone or in combination with $0.25 \times 10^6$ allogeneic T cells.

#### Crypt isolation

Intestinal crypts were isolated as previously described[15] . Briefly, the small intestine (SI) was flushed with PBS, opened longitudinally, and cut into 1 cm segments. Tissue fragments were washed repeatedly until the supernatant was clear. Samples were then incubated in PBS containing 5 mM EDTA at 4°C for 30 min. Crypts were released by vigorous shaking and collected for further processing.

#### Organoid recovery assay

Crypt isolation was performed as described above on day 7 after TBI. Upon isolation, crypts were counted and seeded at defined density (200 crypts/drop for small intestine). Regeneration was assessed by organoid counting after 3 days. SI organoids were cultured in ENR medium.

##### *In vitro* organoid culture

Crypts were embedded in growth factor-reduced Matrigel (Corning) consisting of 66% Matrigel and 33% ENR medium or PBS and plated as 30 µL droplets (~200 crypts/drop) in 24-well plates. Cultures were maintained at 37°C with 5% $CO_2$, and ENR medium was replaced every 2–3 days. Organoids were passaged after 7 days by mechanical dissociation using a 5 mL syringe and an 18G needle. For *in vitro* assays, organoids were used from the first passage onwards.

For murine organoid assays, treatments were added 3 hours after seeding the organoids. Specific treatment conditions are detailed in the corresponding figure legends (H151 = 0.5 µg/mL; IFN-I Ab = 10 µg/ml; IFN-γ = 2.5 ng/ml). Representative brightfield images were acquired on day 6 or after passaging. Regenerative capacity was assessed 3 days after passaging by quantifying organoid numbers relative to control by blinded manual counting or using automated counting with the Incucyte SX5 Live-Cell Analysis System.

Human organoids were derived from large intestinal biopsies and used from passage 3 onwards. Cultures were maintained in WENR medium, with medium changes every other day. Experimental procedures were otherwise identical to murine organoids.

##### Organoid genotoxic damage

Established organoids received treatment of interest (details in figure legends) and were subsequently irradiated (murine: 4 Gy; human: 2 Gy) or subjected to chemotherapeutics agents (cisplatin 10 µM or busulfan 10 µM). Viable organoids were quantified on day 6 post damage induction. Regenerative capacity was assessed as described above.

##### Oxygen consumption

For oxygen consumption measurements of organoids were performed using the PreSens technology (PreSens Precision Sensing GmbH). Organoids were passaged and seeded above the sensor unit of a 24-well Oxodish OD24 plate and cultured for 3 days. Organoids were then treated with H-151 and irradiated after 3h. After the irradiation the media was replaced. 3 h after the last media change, measurements were started for the indicated time frames.

##### Apoptosis in organoid culture

Murine SI established organoids were treated with H151 (0.5 µg/ml) 3 hours prior to irradiation (4 Gy). Apoptosis was quantified three hours after irradiation, by measuring caspase-3/7 activity using the Caspase-Glo® 3/7 Assay (Promega, G8090), according to the manufacturer's instructions.

##### Human organoids

Human organoids were passaged as described above. On day 3 after passaging, organoids were pretreated with G140 at 5 µM, SN-011 at 5 µM, or H151 at 0.5 µg/mL for 3 h before irradiation with

2 Gy. On day 7, organoid number and size were quantified using the Incucyte SX5 Live-Cell Analysis System. To assess regenerative capacity after treatment, organoids were passaged again on day 7 under identical conditions. Organoid outgrowth was monitored, and organoid numbers were quantified on day 5 after passaging.

#### Immunohistochemistry

After deparaffinization and re-hydration, slides were subjected to heat-induced antigen retrieval in a citrate-based buffer, pH 6 for 2.5 min using a steamer at 110°C. Blocking was done with peroxidase blocking solution, Dako REAL (cat no. S2023). Immunohistochemistry (IHC) was performed using the following primary antibodies: CD3 (1:50; clone SP7, catalog no. RM-9107-S1, Thermo Fisher), Ki67 (1:100, Epredia™ Lab Vision™, catalog no. 12683697 , cleaved Caspase-3 (1:1000 Cell Signalling, catalog no. 9661L). Primary antibodies were diluted in Antibody Diluent, Dako REAL (cat no. S2022) and incubated for 30 to 60 min at room temperature. Slides were washed using wash buffer from Dako/Agilent (10x concentrate, cat no. S3006). Secondary antibody (ready to use; for CD3, cat no. 414341F, Histofine; for Ki67, cat no. BA-4001, Vector; for Caspase-3, cat no. BA-1000, Vector cat no. 414341F, Histofine) was incubated for 45 to 60 min at room temperature. Detection was done using the DAB+ substrate detection system (catalog no. K3468, Dako). All stains were validated by internal and/or external positive controls as well as negative control specimens. The slide scanner Pannoramic 1000 Flash RX® (Sysmex Europe SE) was used for image acquisition, choosing the 20x objective. Virtual images from the slide scans were extracted for demonstration purposes using the software CaseViewer version 2.4 (Sysmex Europe SE).

The degree of CD3+ T-cell infiltration was examined in a blinded fashion by a board-certified pathologist. The degree of epithelial T cell infiltration was assessed by counting the number of infiltrating CD3+ cells (intraepithelial lymphocytes) in 10 high power fields (HPF) and calculating the average number of infiltrating cells per HPF for each specimen.

The GvHD pathology was examined in a blinded fashion by a board-certified pathologist (D.H.). Histopathologic scores were assigned based on established criteria[16] .

The number of cleaved Caspase-3+ cells was evaluated as the mean in 10 high-power fields.

The number of Ki67-positive cells was evaluated in individual, well-orientated crypts.

#### Lgr5 in situ hybridization

For detection of Lgr5 expression in FFPE mouse tissue, in situ hybridization (ISH) was performed using RNAscope® 2.5 HD Reagent Kit-BROWN (322300; ACD, Hayward, CA, USA) with the RNAscope® Probe- Mm-Lgr5 (312171; ACD) specific for Lgr5 RNA in mouse. Tissue sections of 4 µm thickness were deparaffinized in xylene, rehydrated in graded ethanol and blocked with peroxidase (10 min). Slides were boiled in kit-provided antigen retrieval buffer at 95 °C for 15 min and digested afterwards with protease at 40 °C for 30 min. For hybridization, tissue sections were incubated with the target probe in the HybEZ hybridization oven (ACD) at 40 °C for 2 h. Pre-amplification and amplification steps were conducted using kit-provided reagents according to the manufacturer's recommendations. For signal detection, sections were incubated with the BROWN

Reagent for 10 min at room temperature followed by counterstaining with hematoxylin, dehydration in ethanol and xylene and mounting with a xylene-based mounting medium.

##### T cell/organoid co-cultures without direct interaction

Murine T cells were isolated from splenocytes of BALB/c mice using the Pan T Cell Isolation Kit II (Miltenyi Biotec, 130-095-130) according to the manufacturer’s instructions. For co-culture, 1 × $10^4$ allogeneic T cells were added to C57BL/6-derived SI organoids in the presence of IL-2 (30 U/mL; PeproTech) and Dynabeads™ Mouse T-Activator CD3/CD28 (Thermo Fisher; 2 µL per well).

To prevent direct contact, plates were maintained at a slight tilt, allowing T cells to settle at the bottom of the well while organoids remained embedded in Matrigel droplets. After 4 days, T cells were removed and medium was replaced with fresh ENR medium. Organoids were passaged on day 7, and regenerative capacity was assessed 3 days later.

##### qPCR and analysis of STING expression in patient GI biopsies

For qPCR analysis, intestinal biopsies were immediately placed in 500 µL RNAlater (QIAGEN) upon collection and stored at −80 °C until processing. Total RNA was extracted using the RNeasy Mini Kit (QIAGEN) according to the manufacturer’s instructions. RNA concentration and integrity were assessed using a NanoDrop spectrophotometer and Bioanalyzer, respectively.

For cDNA synthesis, 1 µg of total RNA was reverse transcribed using Moloney murine leukemia virus reverse transcriptase (Promega) following the manufacturer’s protocol. Quantitative PCR was performed on a Mastercycler Ep Realplex system (Eppendorf) using the QuantiFast SYBR Green PCR Kit (QIAGEN). Gene expression levels were normalized to 18S ribosomal RNA as the internal reference.

Primer sequences used:

STING, forward: 5’-CCT-GAG-TCT-CAG-AAC-AAC-TGC-C-3’, reverse: 5’-GGT-CTT-CAA-GCT-GCC-CAC-AGT-A-3’

18S, forward: 5’-ACC-GAT-TGG-ATG-GTT-TAG-TGA-G-3’, reverse: 5’-CCT-ACG-GAA-ACC-TTG-TTA-CGA-C-3’.

Clinical data were analyzed in R (v4.5.1) using the tidyverse (v2.0.0) suite. Patients were stratified according to STING expression levels (quantitative PCR) measured in GI biopsies, with the cutpoint determined by maximum Youden index using the cutpointr (v1.2.1) package[17]. Patients were also stratified by graft-versus-host disease (GvHD) severity using the Lerner grading system (low, grade <3; high, grade ≥3). Only patients with biopsies obtained within 180 days post-stem cell transplantation were included. Kaplan-Meier survival curves were generated using the ggsurvfit (v1.2.0) and survival (v3.8-6) packages.

##### Media and reagents

Ba/F3 leukemia cells were expanded in RPMI 1640 (Invitrogen) media supplemented with FCS 10% (Gibco), 100 µ/ml penicillin and 100 µg/ml streptomycin (Sigma–Aldrich), β-mercaptoethanol 0.1% (Gibco), L-glutamine 1% (Sigma) and Normocin 0.1% ((InvivoGen).

FLT3-ITD/MLL-PTD AML were reconstituted after thawing in 50% FBS and 50% RPMI 1640.

Human T cell media contained 10% (v/v) FCS, L-glutamine 1% (Sigma), 100 U/ml penicillin and 100 µg/ml streptomycin and was supplemented with 30U/ml hIL-2 (Peprotech). ENR media contained advanced DMEM/F12 (Life technologies), 2 mM L-glutamine (Sigma), 10 mM HEPES (Life technologies), 100 µ/ml penicillin, 100 µg/ml streptomycin (Life technologies), 1.25 mM N-acetyl cysteine (Sigma), 1× B27 supplement (Life technologies), 1× N2 supplement (Life technologies), 50 ng/ml mEGF (Peprotech), 100 ng/ml rec. mNoggin (Peprotech), 10% human R-spondin-1-conditioned medium of hR spondin-1 transfected HEK 293T cells.

#### Statistical analysis

Statistical analyses were performed using GraphPad Prism 10. Unless otherwise specified, data are presented as mean ± SEM. Normality was assessed with the Shapiro–Wilk test. For normally distributed data, comparisons were conducted using two-tailed unpaired Student's *t* test or one-way ANOVA with Tukey's multiple comparisons test. For non-normally distributed data, the Mann–Whitney *U* test or Kruskal–Wallis test with Dunn's post hoc multiple comparisons were applied. Normalized data were analyzed using a one-sample *t* test, with the null hypothesis defined as a control group mean equal to 1. All *in vitro* results were confirmed in at least three independent experiments. Survival analyses (Kaplan–Meier curves) were performed using the log-rank (Mantel–Cox) test. A two-sided significance level of $\alpha = 0.05$ was applied, and the null hypothesis was rejected at $p \leq 0.05$.

#### Bibliography


1. Rückert T, Andrieux G, Boerries M, et al. Human β-defensin 2 ameliorates acute GVHD by limiting ileal neutrophil infiltration and restraining T cell receptor signaling. *Sci Transl Med*. 2022;14(676):eabp9675.

2. Wang Y, Song W, Wang J, et al. Single-cell transcriptome analysis reveals differential nutrient absorption functions in human intestine. *J Exp Med.* 2020;217(2):e20191130.

3. Germain PL, Lun A, Garcia Meixide C, Macnair W, Robinson M. Doublet identification in single-cell sequencing data using scDblFinder. *F1000Res*. 2021;28;10:979.

4. Andreatta M, Carmona SJ. UCell: Robust and scalable single-cell gene signature scoring. *Comput Struct Biotechnol J*. 2021;19:3796-3798.

5. Tirosh I, Izar B, Prakadan SM, et al. Dissecting the multicellular ecosystem of metastatic melanoma by single-cell RNA-seq. *Science*. 2016;352(6282):189-196.

6. Korsunsky I, Millard N, Fan J, et al. Fast, sensitive and accurate integration of single-cell data with Harmony. *Nat Methods*. 2019;16(12):1289-1296.

7. McInnes L, Healy J. UMAP: Uniform Manifold Approximation and Projection for Dimension Reduction. *J. Open Source Softw.* 2018;3(29):861.

8. Shen WK, Zhang CY, Gu YM, et al. An automatic annotation tool and reference database for T cell subtypes and states at single-cell resolution. *Sci Bull (Beijing)*. 2025;70(10):1659-1672.

9. Aran D, Looney AP, Liu L, et al. Reference-based analysis of lung single-cell sequencing reveals a transitional profibrotic macrophage. *Nat Immunol*. 2019;20(2):163-172.

10. Haber AL, Biton M, Rogel N, et al. A single-cell survey of the small intestinal epithelium. *Nature*. 2017;551(7680):333-339.

11. He L, Davila-Velderrain J, Sumida TS, Hafler DA, Kellis M, Kulminski AM. NEBULA is a fast negative binomial mixed model for differential or co-expression analysis of large-scale multi-subject single-cell data. *Commun Biol*. 2021;4(1):629.

12. Subramanian A, Tamayo P, Mootha VK, et al. Gene set enrichment analysis: A knowledge-based approach for interpreting genome-wide expression profiles. *Proc Natl Acad Sci U S A.* 2005;102(43):15545-50.

13. Liberzon A, Birger C, Thorvaldsdóttir H, Ghandi M, Mesirov JP, Tamayo P. The Molecular Signatures Database Hallmark Gene Set Collection. *Cell Syst*. 2015;1(6):417-425.

14. Venables B, Ripley B. Modern Applied Statistics with S. Springer, NY; 2002.

15. Sato T, Vries RG, Snippert HJ, et al. Single Lgr5 stem cells build crypt-villus structures in vitro without a mesenchymal niche. *Nature*. 2009;459(7244):262-265.

16. Lerner KG, Kao GF, Storb R, Buckner CD, Clift RA, Thomas ED. Histopathology of graft-vs.-host reaction (GvHR) in human recipients of marrow from HL-A-matched sibling donors. *Transplant Proc*. 1974;6(4):367-371.

17. Thiele C, Hirschfeld G. cutpointr: Improved Estimation and Validation of Optimal Cutpoints in R. *J Stat Softw*. 2021;98(11):1-27.

#### Tables legend

| Supplemental table 1 | List of antibodies |
|---|---|

| **Supplemental table 2** | |
|---|---|
| (ref to Figure 2C) | Gene Set Enrichment Analysis (GSEA) results for scRNA-seq of human PanT isolated from healthy donor PBMCs. Each worksheet corresponds to a distinct cell population. GSEA was performed on ranked differential gene expression data for the indicated comparison. Enrichment results for pathways from different databases are reported together with nominal p-values, FDR-adjusted p-values, normalized enrichment scores (NES), pathway size, and leading-edge genes contributing to pathway enrichment. |

| **Supplemental table 3** | |
|---|---|
| (ref to Figure 3C) | Gene Set Enrichment Analysis (GSEA) results for scRNA-seq of epithelial cells isolated from the murine intestine. Each worksheet corresponds to a distinct cell population. GSEA was performed on ranked differential gene expression data for the indicated comparison. Differential expression analysis and GSEA have been performed multiple times with different random samples of cells (see methods). Enrichment results for pathways from different databases are reported together with the average of the log FDR-adjusted p-values across random samples and average normalized enrichment scores (NES). |

| **Supplemental table 4** | |
|---|---|
| ref to SF4A, DE ileum | Published scRNA-seq dataset from the human intestine. Differential gene expression analysis of STING and cGAS between cell types in the ileum. |
| ref SF4B, DE colon | Published scRNA-seq dataset from the human intestine. Differential gene expression analysis of STING and cGAS between cell types in the colon. |
| ref to SF4AB, DE between segments | Published scRNA-seq dataset from the human intestine. Differential expression of STING and cGAS between segments of the intestine. |
| ref to SF4F | scRNA-seq of epithelial cells isolated from mice. |

| | |
|---|---|
| | Differential abundance analysis measures changes in the relative abundance of cell populations between H151-treated and control samples. Each row represents a single population |
| SingleR annotation proportions | scRNA-seq of epithelial cells isolated from the murine intestine.<br>SingleR annotation proportions of different cell types among epithelial cells. Each column corresponds to one of the cell type annotations indicated in the UMAP in SF4E, and each row corresponds to the automatic SingleR annotation label |
| “DE stress prog vs Entero all” | Differential gene expression analysis results for scRNA-seq of epithelial cells isolated from the murine intestine. Differential expression analysis was performed to identify marker genes enriched in the “stressed progenitors” population relative to other cell types. This table specifically reports differential expression in comparison to enterocytes-like cells. |
| DE stress prog vs stem | Differential gene expression analysis results for scRNA-seq of epithelial cells isolated from the murine intestine<br>Differential expression analysis was performed to identify marker genes enriched in “stressed progenitors” population relative to other cell types. This table specifically reports differential expression in comparison to the “stem” population. |
| ref to SF4G | Gene Set Enrichment Analysis (GSEA) results for scRNA-seq of epithelial cells isolated from the murine intestine. Here “stressed progenitors” are compared to all other cell types. |

| | |
|---|---|
| **Supplemental table 5** | Summary of patient characteristics |

**Supplemental table 1:** Overview of all utilized antibodies for organoids, flow cytometry and scRNAseq with specific dilutions.

| Antigen | Fluorochrome | RRID | Company | Dilution |
|---|---|---|---|---|
| m IFNAR1 | in vivo Antibody | AB_2687723 | BioXcell | 10 µg/ml |
| m CD25 | BV650 | AB_11125760 | Biolegend | 1:100 |
| m CD45.2 | BUV737 | AB_2870107 | BD | 1:100 |
| m CD326 | BUV395 | AB_2740020 | BD | 1:100 |
| m CD45 | Alexa Fluor 700 | AB_493714; AB_493715 | Biolegend | 1:100 |
| m CD11b | BUV395 | AB_2738276 | BD | 1:300 |
| m CD8a | BUV395 | AB_2732919 | BD | 1:200 |
| m CD366 (Tim3) | BV480 | AB_2744184 | BD | 1:100 |
| m CD3 | FITC | AB_312660; AB_312661 | Biolegend | 1:200 |
| m CD279 (PD-1) | PE | AB_1877232; AB_1877231 | Biolegend | 1:250 |
| m CD4 | PerCP/Cyanine5.5 | AB_893330; AB_893324 | Biolegend | 1:200 |
| m IFN-γ | PE/Cyanine7 | AB_1595591; AB_2295770 | Biolegend | 1:500 |
| m (C57BL) H-2Kb | Brilliant Violet 421 | AB_2876430 | Biolegend | 1:150 |
| m/h FoxP3 | Alexa Fluor 647 | AB_439749; AB_439750 | Biolegend | 1:100 |
| m I-A/I-E | FITC | AB_313320; AB_313321 | Biolegend | 1:1500 |
| m CD11c | PE/Cy7 | AB_493569 | Biolegend | 1:300 |
| m F4/80 | PE/Cyanine7 | AB_2562305 | Biolegend | 1:100 |
| m CD64 | Brilliant Violet 650 | AB_3106221 | Biolegend | 1:100 |
| m CD83 | Brilliant Violet 650 | AB_11203713 | Biolegend | 1:100 |
| m CD86 | PE | AB_313150; AB_313151 | Biolegend | 1:200 |
| m CD80 | PerCP/Cyanine5.5 | AB_893406; AB_2291392 | Biolegend | 1:100 |
| m CD62L | PE/Dazzle 594 | AB_2566162 | Biolegend | 1:100 |
| m CD44 | BV750 | AB_2941373 | Biolegend | 1:100 |
| m CD117 | PE | AB_2734235 | Biolegend | 1:100 |
| m CD32/16 | Fc blocking | AB_312800 | Biolegend | 1:100 |
| h CD45 | APC | AB_314399 | Biolegend | 1:100 |
| h CD25 | BUV737 | AB_2870132 | BD | 1:100 |
| h CD8a | BUV805 | AB_2871326 | BD | 1:100 |
| h PD-1 | Brilliant Violet 421 | AB_2721517 | Biolegend | 1:50 |
| h TIM-3 | Brilliant Violet 605 | AB_2741099 | BD | 1:100 |
| h T-bet | Brilliant Violet 650 | AB_2738616 | BD | 1:100 |
| h CD4 | Brilliant Violet 711 | AB_2737965 | BD | 1:100 |
| h CD3 | Brilliant Violet 785 | AB_11219196 | Biolegend | 1:100 |
| h Perforin | FITC | AB_493252 | Biolegend | 1:50 |
| h IFNγ | BB700 | AB_2744484 | BD | 1:100 |
| h Granzyme B | PE-Dazzle 594 | AB_2728382 | Biolegend | 1:100 |
| Foxp3 | PE-Cy5 | AB_10597134 | Thermo Fisher | 1:100 |
| h TNFα | BUV395 | AB_2738533 | BD | 1:100 |
| viability dye | zombie NIR | | Biolegend | 1:1000 |
| viability dye | zombie UV | | Biolegend | 1:1000 |
| viability dye | Propidium Iodide | | Biolegend | 1:10 |
| viability dye | zombie Aqua | | Biolegend | 1:1000 |
| TotalSeq™-B0301 anti-mouse Hashtag 1 | Oligo Hashtag | AB_2814067 | Biolegend | 1:100 |
| TotalSeq™-B0302 anti-mouse Hashtag 2 | Oligo Hashtag | AB_2814068 | Biolegend | 1:100 |
| TotalSeq™-B0303 anti-mouse Hashtag 3 | Oligo Hashtag | AB_2814069 | Biolegend | 1:100 |
| TotalSeq™-B0304 anti-mouse Hashtag 4 | Oligo Hashtag | AB_2814070 | Biolegend | 1:100 |
| TotalSeq™-B0305 anti-mouse Hashtag 5 | Oligo Hashtag | AB_2814071 | Biolegend | 1:100 |
| TotalSeq™-B0306 anti-mouse Hashtag 6 | Oligo Hashtag | AB_2814072 | Biolegend | 1:100 |

**Supplemental table 5:** Summary of patient characteristics

| **Patients (N=77)** | | | |
|---|---|---|---|
| | | **mean** | **(range)** |
| **Age in years** | | 53 | (20-69) |
| **Sex** | | **n** | **(%)** |
| | male | 46 | 59,7 |
| | female | 31 | 40,3 |
| **Diagnosis** | | | |
| | Acute leukemia | 41 | 53,2 |
| | Myelodysplastic syndrome | 11 | 14,3 |
| | Myeloproliferative syndrome | 3 | 3,9 |
| | Lymphoma | 22 | 28,6 |
| **Stage of underlying disease** | | | |
| | early | 16 | 20,8 |
| | intermediate | 27 | 35,1 |
| | advanced | 34 | 44,2 |
| **Donor type** | | | |
| | Unrelated donor | 49 | 63,6 |
| | Sibling | 25 | 32,5 |
| | Haploidentical donor | 3 | 3,9 |
| **Stem cell source** | | | |
| | PBSC | 71 | 92,2 |
| | BM | 6 | 7,8 |
| **Conditioning regimen** | | | |
| | Reduced intensity | 69 | 89,6 |
| | Standard | 8 | 10,4 |
| **Immunosuppression** | | | |
| | CyA/MTX | 58 | 75,3 |
| | CyA/MMF | 14 | 18,2 |
| | Everolimus | 1 | 1,3 |
| | Tacro/MMF | 1 | 1,3 |
| | PostTxCy/Tacro/MMF | 3 | 3,9 |
| **Clinical GvHD overall grade max.** | | | |
| | no GvHD | 25 | 32,5 |
| | Grade I | 9 | 11,7 |
| | Grade II | 21 | 27,3 |
| | Grade III | 11 | 14,3 |
| | Grade IV | 11 | 14,3 |
| **Lerner GvHD** | | | |
| | Lerner 0 | 28 | 36,4 |
| | Lerner 1 | 26 | 33,8 |
| | Lerner 2 | 11 | 14,3 |
| | Lerner 3 | 9 | 11,7 |
| | Lerner 4 | 3 | 3,9 |

### Supplemental figure 1

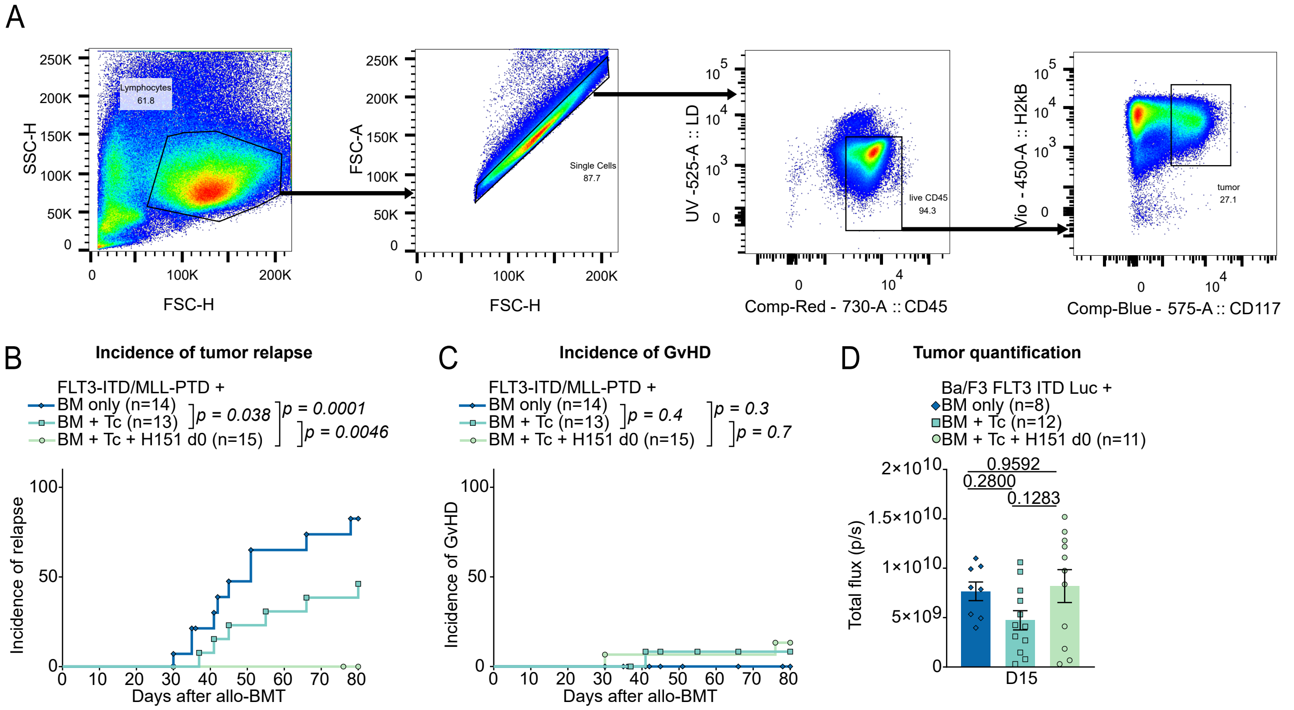

**Supplemental figure 1. Additional data related to Figure 1**

**(A)** Gating strategy used for FACS analyses of leukemia cells in GvL-MLL model. Incidence of relapse **(B)** or GvHD **(C)** of mice following tumor transplantation and allo-BMT. Statistics by Log-rank test. **(D)** Quantification of BLI signal on day 15 to monitor leukemia progression. Data were pooled from two independent experiments. P values obtained by one-way ANOVA with Tukey's multiple comparisons test. All data are shown as mean ± SEM.

### Supplemental figure 2

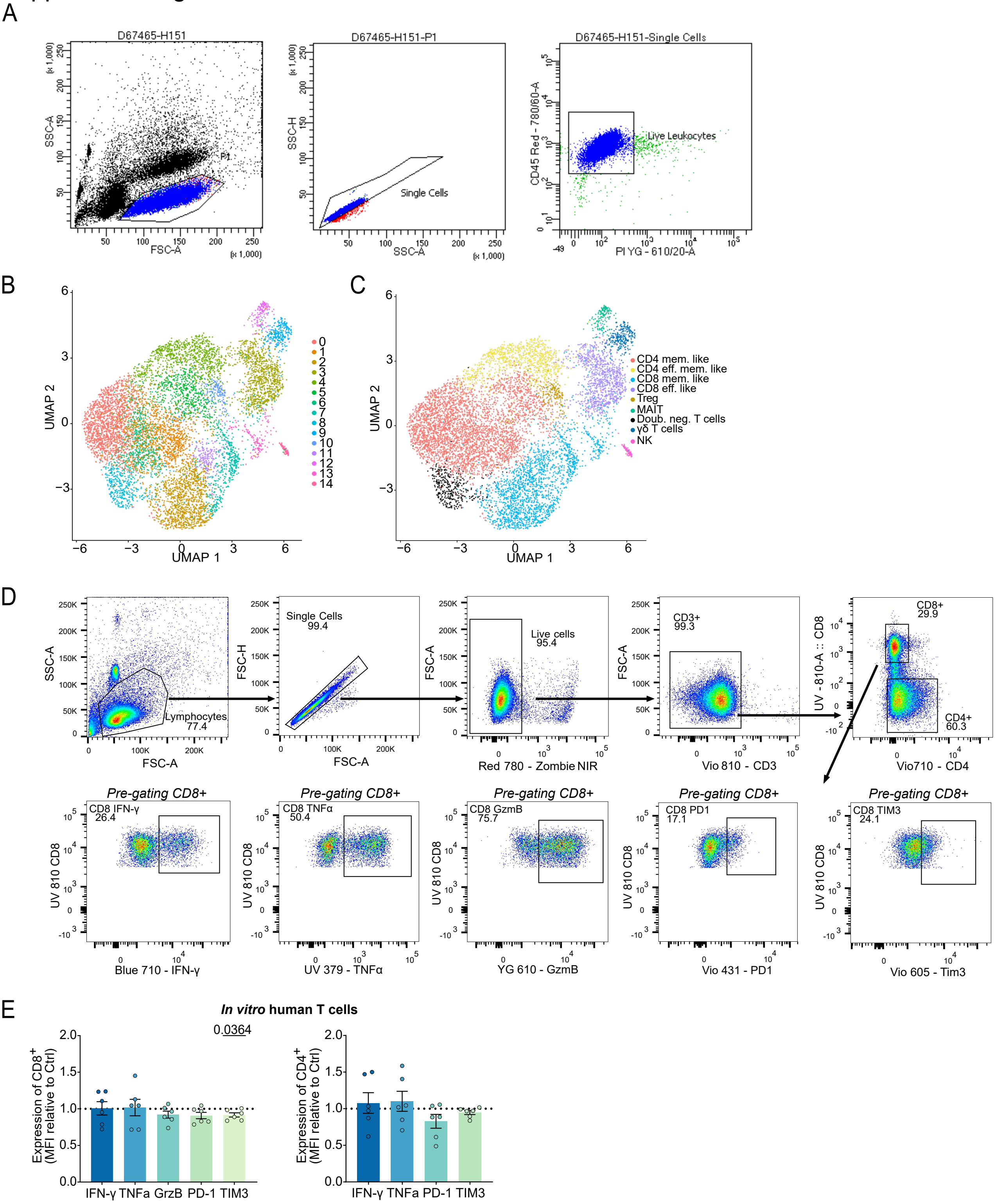

**Supplemental figure 2. Additional data related to Figure 2**

**(A)** Gating strategy for sorting of live CD45+ cells from the cell suspension of Pan T cells from healthy donors. **(B)** UMAP plot of human pan-T scRNA-seq data colored by graph-based clustering, across all experimental groups. **(C) )** UMAP plot of human pan-T scRNA-seq colored by cell type annotation, across all experimental groups. **(D)** Gating strategy for FACS analysis of human pan-T cells isolated from healthy donors, stimulated with anti-CD3/CD28 beads, 30 U/mL human IL-2, and H151 for 24 h prior to flow cytometric analysis. **(E)** Protein expression levels (MFI) in CD8+ and CD4+ cells, normalized to the respective control. Samples were pooled from six healthy donors and were analyzed using a one sample t-test.

Supplemental figure 3

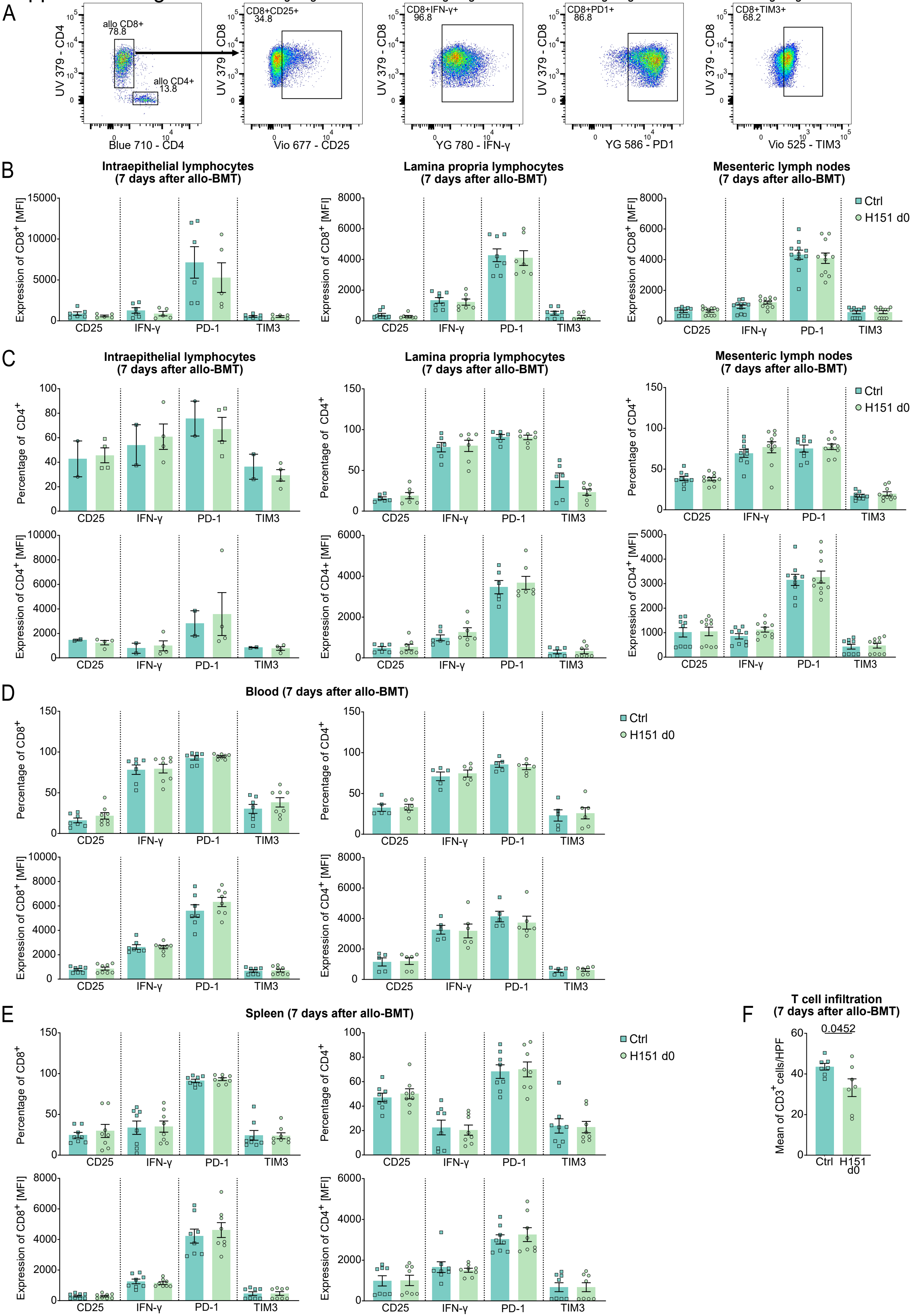

**Supplemental figure 3. Additional data related to Figure 2**

**(A)** Gating strategy used for FACS analysis of T cells isolated from different mouse organs at day 7 after allo-BMT. $CD8^+$ and $CD4^+$ cells were gated on live allogeneic $CD3^+$ cells. **(B)** Mean of fluorescence intensity (MFI) of activation markers (CD25, IFN-γ, PD-1, and TIM3) expressed by $CD8^+$ T cells derived from intraepithelial lymphocytes (IEL), lamina propria (LP), and mesenteric lymph nodes (mLN). **(C)** Frequency of $CD4^+$ T cells positive for the indicated markers and corresponding expression levels, reported as mean fluorescence intensity (MFI). **(D)** Frequency and expression levels of the indicated markers in $CD8^+$ and $CD4^+$ T cells isolated from peripheral blood at day 7 after allo-BMT. **(E)** Frequency and expression levels of the indicated markers in $CD8^+$ and $CD4^+$ T cells isolated from spleen at day 7 after allo-BMT. **(F)** Counts of infiltrating T cells in the mice ileum at day 7 after allo-BMT. Data were pooled from 3 independent experiments (n = 7 (Ctrl) or 7 (H151 d0)). Statistical analysis by unpaired t-test. All data are shown as mean ± SEM.

Supplemental figure 4

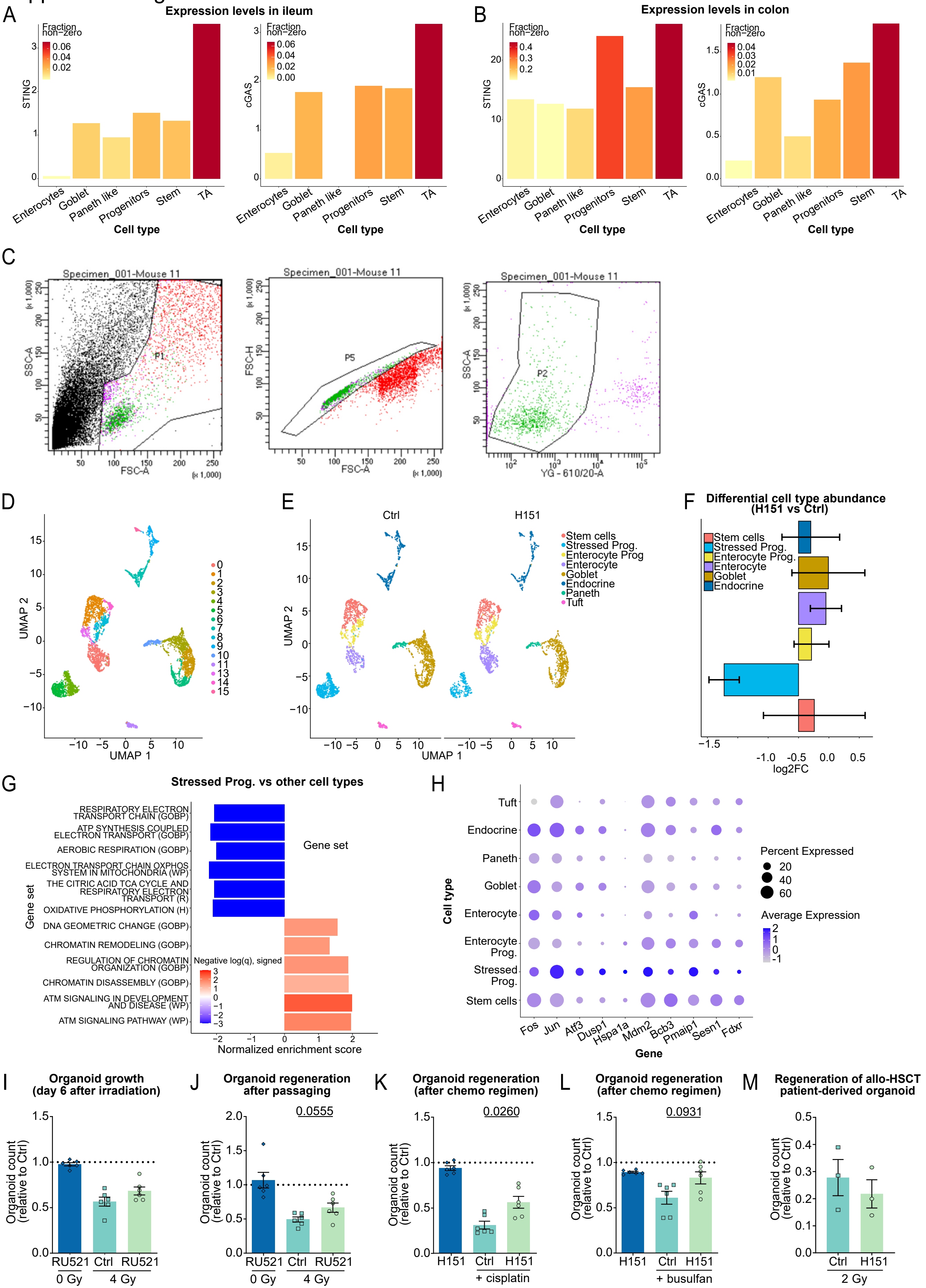

**Supplemental figure 4. Additional data related to Figure 3**

Bar plot of STING and cGAS gene expression in different cell types across **(A)** ileum and **(B)** colon, from a published scRNA-seq dataset of the human intestine. Gene expression is shown as aggregated UMI counts across all single cells of a given cell type (pseudobulk), per intestinal segment. Normalization by total pseudobulk UMI counts. The color of the bars corresponds to the fraction of cells with non-zero counts within a given cell type and intestinal segment. **(C)** Gating strategy for sorting of live cells from the cell suspension of intestinal crypts. **(D)** UMAP plot of murine intestinal scRNA-seq data colored by graph-based clustering, across all experimental groups. **(E)** UMAP visualization of single cells colored by final cell type annotation, comparing control and H151-treated groups. **(F)** Differential cell type abundance analysis on cell type counts derived from murine intestinal scRNA-seq data. H151-treated mice are compared to controls using negative binomial regression. Data are presented as $\log_2$ fold-change estimates with corresponding 95% confidence intervals based on a normal distribution. **(G)** Bar plot showing gene set enrichment analysis (GSEA) results for selected pathways in stressed progenitor cells compared to other all other cell types from the murine intestinal scRNA-seq data. Bar color represents the −log10 of the GSEA q-value (FDR), with the sign indicating the direction of regulation (positive = upregulated, negative = downregulated). The scale is capped at -3/3 for better interpretability. Bar length corresponds to the GSEA normalized enrichment score. Gene sets/pathways are derived from the Hallmark (H), Reactome (R), GO Biological Process (GOBP) and WikiPathways (WP) collections of MSigDB. **(H)** Dotmap of selected marker genes related to stress responses across intestinal epithelial cell populations. Columns represent genes and rows represent annotated cell types. Color intensity corresponds to library-size normalized UMI counts, averaged by cell type, logtransformed and scaled across cell types. Dot size corresponds to the percentage of cells with non-zero counts withing a given cell type. Organoid counts were assessed prior to passaging **(I)** and after passaging **(J)**, under steady-state conditions or following 4 Gy irradiation, with or without RU521 treatment. Data were normalized to control (dotted line) and pooled from three independent experiments (n=6 mice). Statistical analysis was performed using unpaired t test with Welch's correction. Organoid numbers were quantified after passaging following treatment with the chemotherapeutic agents cisplatin **(K)** or busulfan **(L)**, with or without H151 administered 3 hours prior to drug exposure. Data were normalized to the control (dotted line) and pooled from three independent experiments (n=6 mice). P values were calculated by Mann Whitney test. **(M)** Growth of human large intestine organoids derived from allo-HSCT patients, treated with the STING inhibitor H151, with 2 Gy irradiation. Data were pooled from three donors across three independent experiments. All data are shown as mean ± SEM.

Supplemental figure 5

A

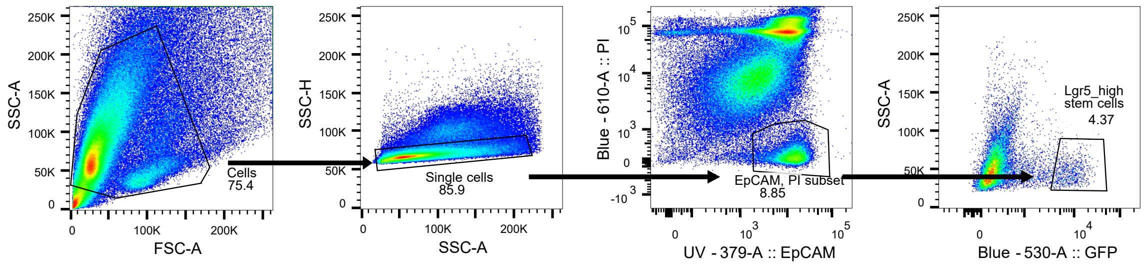

**Supplemental figure 5. Additional data related to Figure 5**

**(A)** Gating strategy used for FACS analysis of epithelial cells derived from murine SI organoids.
